# Metagenomics vs. total RNA sequencing: most accurate data-processing tools, microbial identification accuracy, and implications for freshwater assessments

**DOI:** 10.1101/2022.06.03.494701

**Authors:** Christopher A. Hempel, Natalie Wright, Julia Harvie, Jose S. Hleap, Sarah J. Adamowicz, Dirk Steinke

**Affiliations:** Department of Integrative Biology, University of Guelph, Guelph, ON N1G 2W1, Canada; SHARCNET, University of Guelph, Guelph, ON N1G 2W1, Canada

## Abstract

Metagenomics and total RNA sequencing (total RNA-Seq) have the potential to improve the taxonomic identification of diverse microbial communities, which could allow for the incorporation of microbes into routine freshwater assessments. However, these targeted-PCR-free techniques require more testing and optimization. In this study, we processed metagenomics and total RNA-Seq data from a commercially available microbial mock community using 768 data-processing workflows, identified the most accurate data-processing tools, and compared their microbial identification accuracy at equal and increasing sequencing depths. The accuracy of data-processing tools substantially varied among replicates. Total RNA-Seq was more accurate than metagenomics at equal sequencing depths and even at sequencing depths almost one order of magnitude lower than those of metagenomics. We show that while data-processing tools require further exploration, total RNA-Seq might be a favorable alternative to metagenomics for targeted-PCR-free taxonomic identifications of microbial communities and might enable a substantial reduction in sequencing costs while maintaining accuracy. This could be particularly an advantage for routine freshwater assessments, which require cost-effective yet accurate methods, and might allow for the incorporation of microbes into freshwater assessments.

## INTRODUCTION

Ecosystems are globally deteriorating at an unprecedented speed, causing a rapid biodiversity decline (IPBES, 2019; Pettorelli et al., 2021; WWF, 2020), which negatively affects ecosystem services. Under certain future land use and management scenarios, the future value of global ecosystem services is estimated to decline by up to 51 trillion $US/year until 2050 (Kubiszewski et al., 2017). Consequently, ecosystem protection is gaining increased attention, even on the political scale (IPBES, 2019). Freshwater ecosystems are particularly vulnerable and strongly affected by climate change, pollution, habitat fragmentation, and the introduction of invasive species. As a result, they exhibit some of the highest rates of species loss, placing them among the most threatened ecosystems (Reid et al., 2019). Clearly, there is an urgent need to protect, preserve, and restore freshwater ecosystems, but to do so, we first need to determine their natural status. This is usually achieved by using ecological assessments, which include the assembly of biodiversity inventories. Such inventories can be screened for the presence and abundance of taxa that represent specific environmental conditions, so-called bioindicators (Burger, 2006). Common bioindicators are animals, plants, and diatoms (Bellinger and Sigee, 2015; Haury et al., 2006; Karr, 1981; Resh and Unzicker, 1975); however, recently it has been suggested to include more microbes (prokaryotes and unicellular eukaryotes) because they respond faster to environmental changes (Cordier et al., 2019; Pawlowski et al., 2016; Sagova-Mareckova et al., 2021; Smith et al., 2015).

Biodiversity inventories have been traditionally assembled by assessing the morphology of organisms. However, morphological identification can be biased (Stein et al., 2014; Sweeney et al., 2011) or is often not feasible due to a lack of diagnostic traits (Pawlowski et al., 2012; Will and Rubinoff, 2004). A solution to these issues was the development of DNA-based approaches to identify organisms by using target primers that amplify standardized genetic markers (amplicons), which can be applied to assemble community inventories (Clarridge, 2004; Cristescu, 2014; Janda and Abbott, 2007; Petti et al., 2005; Taberlet et al., 2012). However, this approach comes with its own set of biases, mostly due to varying primer-binding affinities (Alberdi et al., 2018; Krehenwinkel et al., 2017; Piper et al., 2019; Stat et al., 2017).

Alternative approaches for taxonomic identification of communities are metagenomics and metatranscriptomics, which involve shotgun sequencing, i.e., the random fragmentation and sequencing of all DNA or RNA in a sample (Almeida and De Martinis, 2019; Bashiardes et al., 2016; Shakya et al., 2019; Wooley et al., 2010). Metatranscriptomics usually involves a messenger RNA (mRNA) enrichment step to analyze gene expression patterns; however, it is possible to skip the mRNA enrichment step to utilize total RNA for taxonomic identification, including ribosomal RNA (rRNA). This approach has been referred to as double-RNA approach (Urich et al., 2008), metatranscriptomics analysis of total rRNA (Turner et al., 2013), total RNA sequencing (total RNA-Seq) (Bang-Andreasen et al., 2020; Li et al., 2016; Li and Guan, 2017), total RNA-based metatranscriptomics, or total RNA-seq-based metatranscriptomics (Li and Guan, 2017). To distinguish this approach from regular metatranscriptomics, we will use the term *total RNA-Seq* in the balance of this paper. Metagenomics and total RNA-Seq remove the need for morphology or barcode-based sequencing with all their associated biases. Furthermore, they allow for the exploration of functional diversity, which can deliver further information about the status of freshwater ecosystems (Cordier et al., 2020b; Leese et al., 2018).

It has been discussed that for taxonomic identification, total RNA-Seq might have an advantage over metagenomics, especially when it comes to the active part of a community, as total RNA-Seq utilizes active transcripts (Bang-Andreasen et al., 2020; Geisen et al., 2015; Gomez- Silvan et al., 2018). Metagenomics, on the other hand, targets all DNA in a sample, and this also includes the DNA of dead and inactive cells and extracellular DNA, which can make up 40–90% of the total DNA pool (Carini et al., 2016; Torti et al., 2015). Consequently, total RNA-Seq might generate more relevant information for ecological assessments given that it targets the portion of the community that is actively interacting with the environment rather than the entire available genetic material of a community.

Another advantage of total RNA-Seq is that it enriches sequencing data for widely used standard genetic markers because 80–98% of RNA consists of rRNA (Peano et al., 2013; Westermann et al., 2012), including both the small subunit (SSU) and the large subunit (LSU) rRNA markers for prokaryotes (16S and 23S rRNA) and unicellular eukaryotes (18S and 28S rRNA). These can, therefore, make up 37–71% of total RNA-Seq reads (Elekwachi et al., 2017; Yu and Zhang, 2012). In contrast, metagenomics targets all DNA, including non-functional genes, repetitive regions, and genes that are functionally important but contain little taxonomic information. Consequently, SSU and LSU rRNA markers can make up as little as 0.05–1.4% of metagenomics reads (Logares et al., 2014; Yilmaz et al., 2011). Since reference databases to date do not contain the full genomes of most microbial taxa but instead only SSU and LSU rRNA sequences, the substantially higher portion of SSU and LSU rRNA markers in total RNA-Seq reads should give total RNA-Seq an advantage over metagenomics in recovering taxonomically informative sequences at comparable sequencing depths.

To date, several studies compared the taxonomic composition of environmental microbial communities obtained through total RNA-Seq and metabarcoding (Lanzén et al., 2011; Yan et al., 2018) or metagenomics (Lanzén et al., 2011; Shi et al., 2011; Urich et al., 2014). However, a controlled, mock community-based comparison of total RNA-Seq and metagenomics for taxonomic identification of microbial communities is lacking. This also includes the comparison and establishment of data-processing workflows, as results based on High-Throughput Sequencing (HTS) are heavily influenced by the tools used to process the data (Bashiardes et al., 2016; Knight et al., 2018; McIntyre et al., 2017; Quince et al., 2017; Shakya et al., 2019; Vollmers et al., 2017). Such a mock community-based comparison is important because it can reveal implications for the assembly of biodiversity inventories.

For this study, we applied metagenomics and total RNA-Seq to a commercially available microbial mock community consisting of eight bacterial and two eukaryotic species with log-distributed abundances. We tested the central idea that total RNA-Seq recovers more taxonomically informative sequences than metagenomics. Therefore, we evaluated the impact of 768 data-processing workflows on taxonomic identification accuracy at species and genus level and based on abundance and presence-absence (P-A) data for each sequencing method. Then, we determined the most accurate workflow for each sequencing method and evaluation level and compared the accuracy of both sequencing methods at equal sequencing depth to determine the more accurate sequencing method. Furthermore, we investigated the relationship between sequencing depth and accuracy for both sequencing methods. Our aim was to answer the following questions: 1) which is the most accurate data-processing workflow for total RNA-Seq, and does it coincide with the most accurate data-processing workflow for metagenomics? 2) Does total RNA-Seq provide more accurate taxonomic identifications than metagenomics at equal sequencing depth? 3) Does the accuracy of total RNA-Seq increase faster than that of metagenomics with increasing sequencing depth?

## MATERIAL AND METHODS

The overall study design is shown in Figure 1, and further details are given in the following.

**Figure 1:**
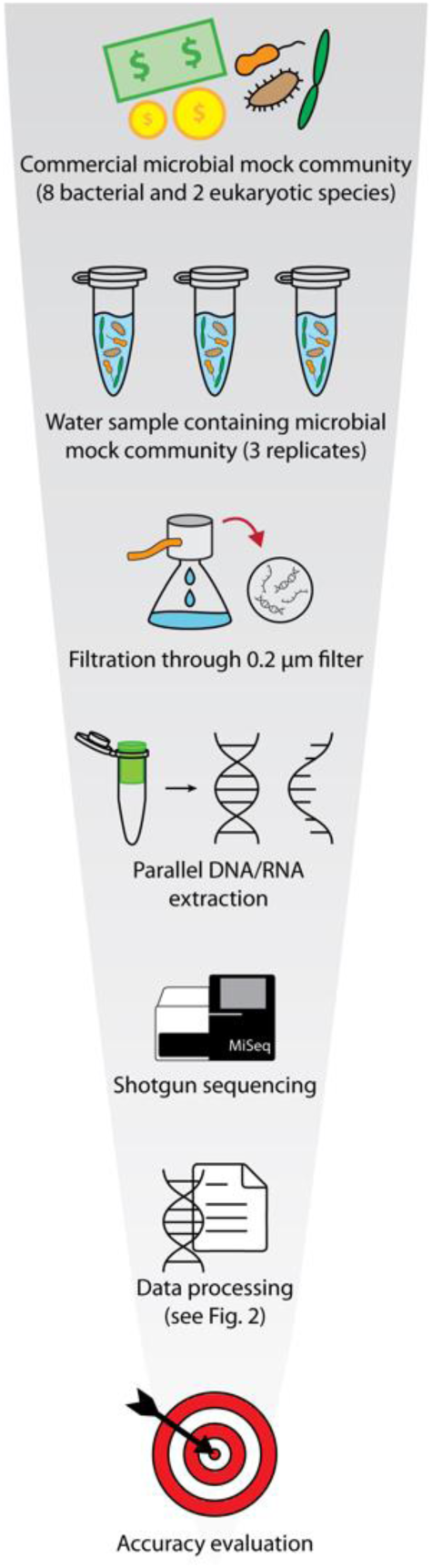
Summary of the study design. Three mock community replicates were obtained by mixing a commercially available microbial mock community with ultrapure water. Samples were filtered through 0.2 µm filters. DNA and total RNA were extracted in parallel and shotgun-sequenced, representing two sequencing methods (metagenomics and total RNA-Seq). The sequencing data were processed using 768 combinations of common data-processing tools, i.e., data-processing workflows. The accuracy of each workflow was statistically evaluated by calculating the Euclidean distance between accuracy metrics determined for each workflow and reference accuracy metrics based on the known mock community composition.

### Microbial mock community

We used a commercially available microbial mock community (ZymoBIOMICS Microbial Community Standard II (Log Distribution); Zymo Research; Irvine; CA U.S.A.), consisting of eight bacterial species (three gram-negative and five gram-positive) and two yeast species with log-distributed species abundances determined by genomic DNA quantity (Table 1). The mock community was preserved in DNA/RNA Shield (Zymo Research; Irvine; CA U.S.A.) to inactivate cells while preserving DNA and RNA. We generated three simulated water sample replicates by adding 130 µl of the mock community containing approximately 381 ng of total DNA to 50 mL ultrapure water respectively.

**Table 1:**
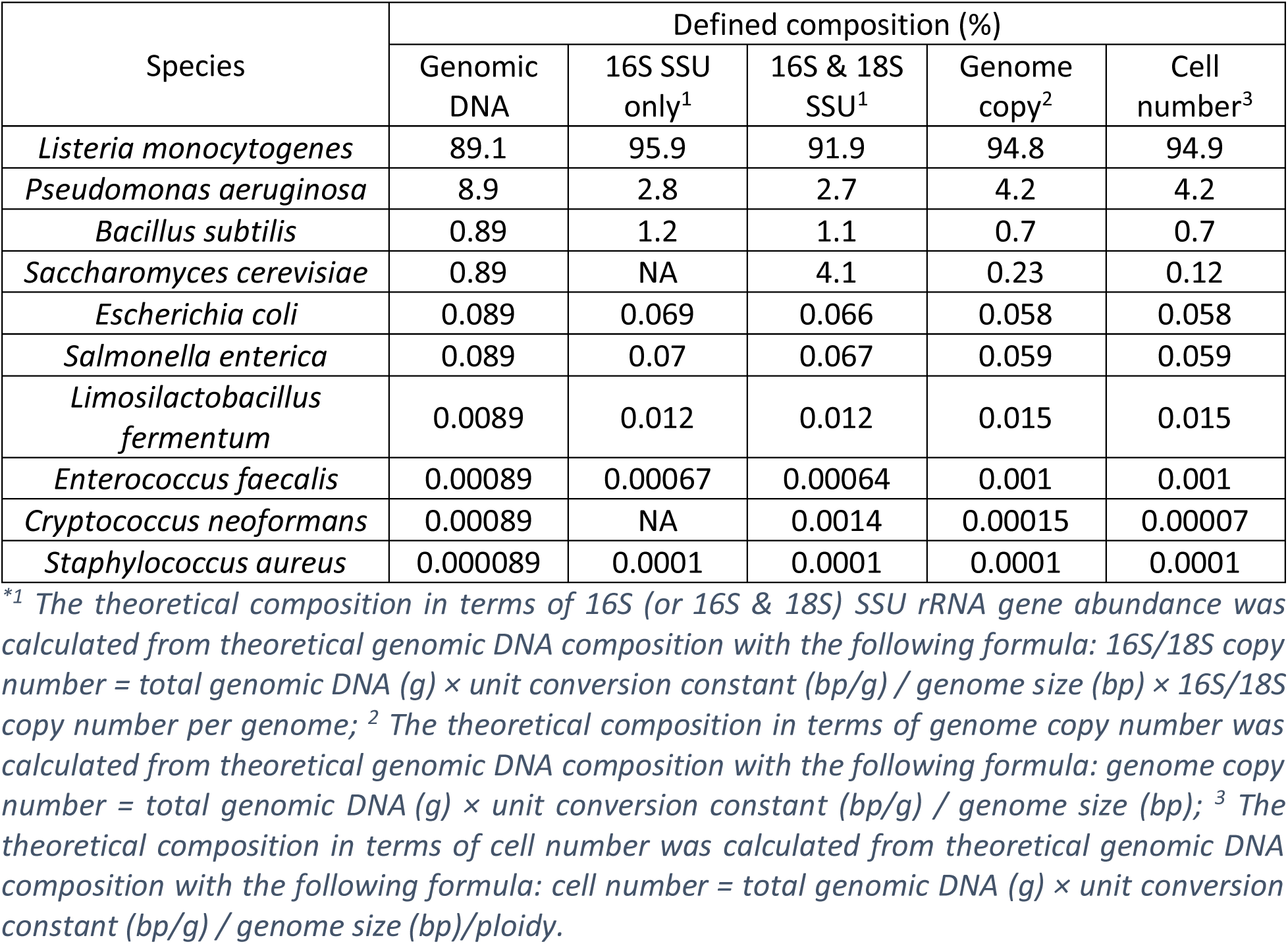
Microbial composition of the mock community (modified from the ZymoBIOMICS Microbial Community Standard II (Log Distribution) manual, for detailed information see manual).

| Species | Defined composition (%) |  |  |  |  |
| --- | --- | --- | --- | --- | --- |
|  | Genomic DNA | 16S SSU only<sup>1</sup> | 16S & 18S SSU<sup>1</sup> | Genome copy<sup>2</sup> | Cell number<sup>3</sup> |
| *Listeria monocytogenes* | 89.1 | 95.9 | 91.9 | 94.8 | 94.9 |
| *Pseudomonas aeruginosa* | 8.9 | 2.8 | 2.7 | 4.2 | 4.2 |
| *Bacillus subtilis* | 0.89 | 1.2 | 1.1 | 0.7 | 0.7 |
| *Saccharomyces cerevisiae* | 0.89 | NA | 4.1 | 0.23 | 0.12 |
| *Escherichia coli* | 0.089 | 0.069 | 0.066 | 0.058 | 0.058 |
| *Salmonella enterica* | 0.089 | 0.07 | 0.067 | 0.059 | 0.059 |
| *Limosilactobacillus fermentum* | 0.0089 | 0.012 | 0.012 | 0.015 | 0.015 |
| *Enterococcus faecalis* | 0.00089 | 0.00067 | 0.00064 | 0.001 | 0.001 |
| *Cryptococcus neoformans* | 0.00089 | NA | 0.0014 | 0.00015 | 0.00007 |
| *Staphylococcus aureus* | 0.000089 | 0.0001 | 0.0001 | 0.0001 | 0.0001 |

<sup>\*1</sup> The theoretical composition in terms of 16S (or 16S & 18S) SSU rRNA gene abundance was calculated from theoretical genomic DNA composition with the following formula: 16S/18S copy number = total genomic DNA (g) × unit conversion constant (bp/g) / genome size (bp) × 16S/18S copy number per genome; <sup>2</sup> The theoretical composition in terms of genome copy number was calculated from theoretical genomic DNA composition with the following formula: genome copy number = total genomic DNA (g) × unit conversion constant (bp/g) / genome size (bp); <sup>3</sup> The theoretical composition in terms of cell number was calculated from theoretical genomic DNA composition with the following formula: cell number = total genomic DNA (g) × unit conversion constant (bp/g) / genome size (bp)/ploidy.

### Laboratory processing

We processed water samples in a separate clean laboratory (for details on handling see dx.doi.org/10.17504/protocols.io.eq2lyn6zrvx9/v1). All samples were filtered through sterile 0.2 µm Nalgene Analytical Test Filter Funnels (Thermo Fisher Scientific; Burlington; ON Canada) using an 80 mbar Welch WOB-L® Dry Vacuum Pump (VWR International; Mississauga; ON Canada). We filtered the three 50 mL microbial mock community mixtures and added a negative filtration control by additionally filtering 50 mL of the ultrapure water that was used to set up the mixtures. After each filtration, we handled the filters with ethanol-, bleach-, and heat-sterilized equipment, immediately cut filters into small pieces, and transferred them into ZR BashingBead Lysis Tubes (0.1 & 0.5 mm) (Zymo Research; Irvine; CA U.S.A.), which were prepared with 1 mL of DNA/RNA Shield under a clean hood in a low DNA-concentration laboratory before filtration.

BashingBead tubes were shaken following the manufacturer’s instructions for optimal cell breakup of the purchased mock community by using a Vortex-Genie 2 (Scientific Industries, Inc.; Burlington; NY U.S.A.) with a Horizontal-(24) Microtube holder (Scientific Industries, Inc.; Burlington; NY U.S.A.) for 40 min at maximum rpm to break up cells.

For parallel DNA and total RNA extraction from samples, we used a modified version of the Quick-DNA/RNA Microprep Plus Kit (Zymo Research; Irvine; CA U.S.A.). We added a purification step using Zymo-Spin II-µHRC Filters (Zymo Research; Irvine; CA U.S.A.) and modified the protocol to process more lysate volume (for the modified protocol see dx.doi.org/10.17504/protocols.io.14egn79bpv5d/v1). We extracted the samples and negative filtration control under a clean hood in a low DNA-concentration laboratory and added a negative extraction control by processing only the extraction buffer along with the other samples.

The extracted nucleic acids and negative filtration and extraction controls were sent to Génome Québec (Montreal; QC Canada) for library preparation and shotgun sequencing on an Illumina MiSeq platform. Processing steps and quality control of DNA and RNA samples are described in Supplemental Material 1. DNA and total RNA concentrations of samples ranged from 2.36–3.05 ng/µL (DNA) and 1.87–3.32 ng/µL (RNA) in 100 µL eluates, while filtration and extraction controls did not contain any DNA but low amounts of RNA (Tables S1 and S2). RNA integrity numbers (RINs) of samples were all N/A (Table S3); however, because the samples contained both prokaryotic and eukaryotic RNA, RINs were not applicable and N/As were ignored. During library preparation, normalization was performed by processing equal volumes of samples instead of equal concentrations of samples. We chose this alternative normalization method because it allowed for an equal relative sequencing depth per sample as opposed to an equal total sequencing depth. That way, the relative number of reads per sample mirrored the relative amount of DNA/RNA, avoiding an over- or underrepresentation of samples with higher or lower DNA/RNA amounts.

### Bioinformatic processing

Sequence processing was divided into five steps: trimming and quality filtering, rRNA sorting, assembly, mapping, and taxonomic annotation (Figure 2). We generated a command-line-based script to run all 768 data-processing workflows, including the processing of each result into a table containing assembled scaffolds, their taxonomic annotations, and their mean depth of coverage. The full code, including scripts to translate SILVA taxonomy into Genbank taxonomy, to create the SILVA kraken2 and BLAST databases, and to filter BLAST results based on CREST and BASTA, is available on GitHub (https://github.com/hempelc/metagenomics-vs-totalRNASeq). All 768 workflows were applied to both metagenomics and total RNA-Seq data and run in parallel using the high-performance computing clusters of Compute Canada.

**Figure 2:**
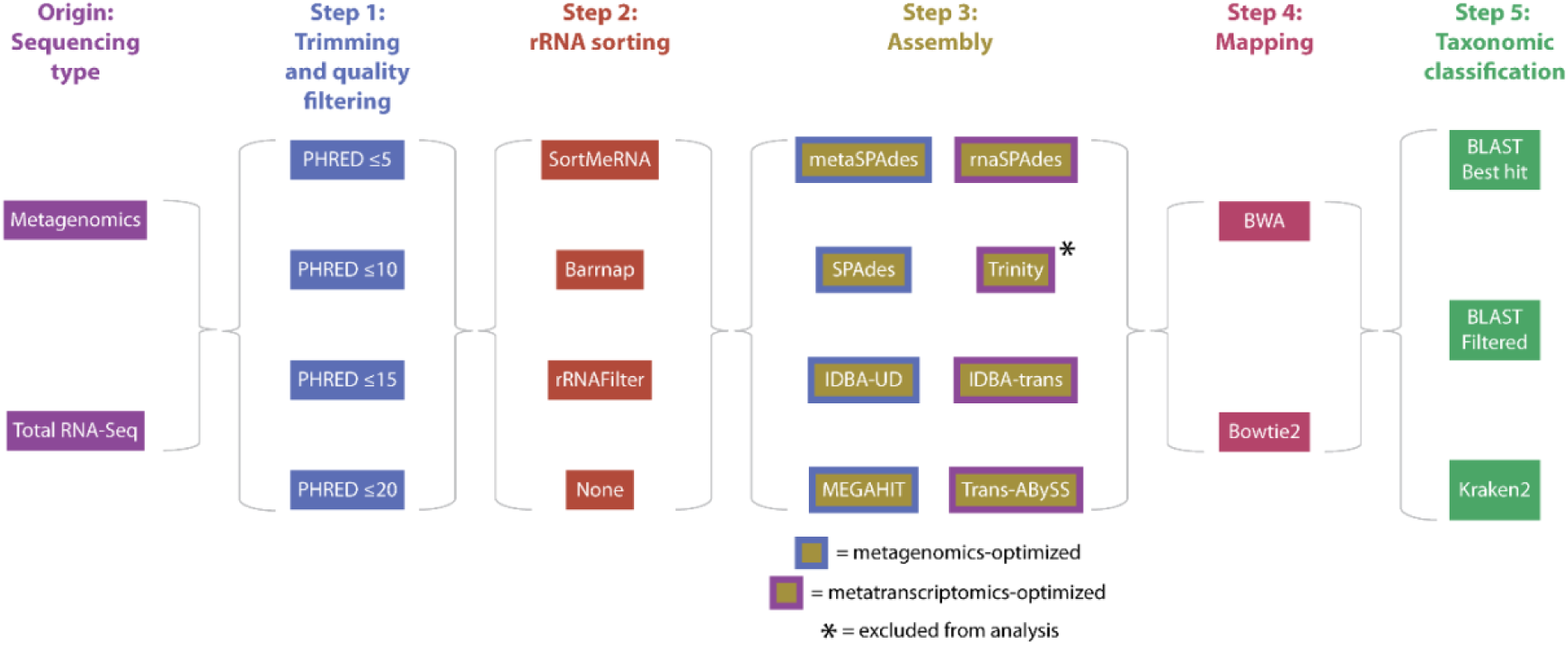
Summary of the 786 workflows applied to metagenomics and total RNA-Seq data, including the steps and tools used to process the data. Note that in step 3, some assemblers are metagenomics- and some metatranscriptomics-optimized, yet we tested all on both metagenomics and total RNA-Seq data. We were unable to run Trinity successfully and excluded it from further analysis (for more details see the methods section “Step three (assembly)”).

#### Step one (trimming and quality filtering)

Recommended PHRED score cut-offs for trimming and quality filtering of HTS data vary across the literature. While strict quality trimming, i.e., trimming at high PHRED score cut-offs of 20–30 is common (Deiner et al., 2017; MacManes, 2014), gentle quality trimming at PHRED score cut-offs of 2–5 can result in better transcript discovery (MacManes, 2014). To explore the effect of different PHRED score cut-offs on both metagenomics- and total RNA-Seq-based taxonomic identification, we used Trimmomatic v0.39 (Bolger et al., 2014) at four different PHRED score cut- offs (PHRED ≤5, ≤10, ≤15, and ≤20). We trimmed the leading and trailing low-quality nucleotides of each read and ran a sliding window of size 4 over each read, cutting if the average quality of nucleotides in the sliding window was below the respective PHRED score cut-offs. After trimming, we excluded reads shorter than 25 nucleotides and error-corrected reads using the error-correction module of the assembler SPAdes v3.14.1 (Bankevich et al., 2012) by running SPAdes on forward and reverse reads with the parameter `--only-error-correction`. That way, error correction between all assemblers was standardized.

#### Step two (rRNA sorting)

We used three approaches to sort reads into rRNA and non-rRNA reads:

(1) alignment-based with SortMeRNA v4.0.0 (Kopylova et al., 2012), which sorts reads by aligning them to built-in reference rRNA databases. Trimmed forward and reverse reads were aligned against all built-in reference rRNA databases using the parameters `-fastx` to generate output files in fasta format, `-num_alignments 1` to only filter the reads, `--paired_in` to keep both forward and reverse reads if only one matched, `--out2` to save forward and reverse reads in separate files, and all other parameters set to default.
(2) Hidden Markov model-based (HMM-based) with barrnap v0.9 (Seemann, unpublished, https://github.com/tseemann/barrnap, accessed on 18 Jun 2021), which predicts the location of rRNA genes in genomes using pre-trained HMMs. However, since barrnap only keeps reads that contain rRNA genes, we used it in this study to only identify rRNA reads. Both trimmed forward and reverse reads were separately run against HMMs for all three domains of life, setting all parameters to default values except the parameters `--lencutoff` and `--reject`, which were set to 0.000001 to keep all partial matches. To keep paired reads, the read names of both filtered forward and reverse reads were concatenated and extracted from the original trimmed forward and reverse reads using Seqtk v1.3-r106 (Li, unpublished, https://github.com/lh3/seqtk, accessed via the Linux package manager).
(3) kmer-based with rRNAFilter v1.1 (Wang et al., 2017). rRNAFilter considers the much higher abundance of rRNA reads compared to non-rRNA reads and filters them based on their k-mer frequencies. Both trimmed forward and reverse reads were separately filtered with default parameters including a k-mer length of 20. To keep paired reads, the read names of both filtered forward and reverse reads were concatenated and extracted from both the original trimmed forward and reverse reads using Seqtk.

For each of the three approaches, non-rRNA reads were subsequently excluded. Additionally, we performed an *unsorted* approach using all reads, leading to four rRNA sorting methods in total.

#### Step three (assembly)

We tested eight assemblers for both metagenomics and total RNA-Seq reads: SPAdes, metaSPAdes v3.14.1 (Nurk et al., 2017), MEGAHIT v1.2.9 (Li et al., 2015), IDBA-UD v1.1.1 (Peng et al., 2012), Trinity v2.10.0 (Grabherr et al., 2013), rnaSPAdes v3.14.1 (Bushmanova et al., 2019), IDBA-tran v1.1.1 (Peng et al., 2013), and Trans-ABySS v2.0.1 (Robertson et al., 2010). Although four of the assemblers are commonly used for metagenome assemblies (SPAdes, metaSPAdes, MEGAHIT, and IDBA-UD) and four for metatranscriptome assemblies (Trinity, rnaSPAdes, IDBA-tran, and Trans-ABySS), we tested them all for both data sets because the read composition of total RNA-Seq data is different from traditional metatranscriptomics, and we wanted to test how both assembler types deal with such an uncommon read composition. All assemblers were run with default parameters except MEGAHIT, for which the parameter `--presets meta-large` was used to adjust the k-mer sizes to better assemble large and complex metagenomes. All assemblers but Trans-ABySS run multiple k-mer lengths by default, whereas Trans-ABySS runs with only one k-mer length of 32 by default. Trans-ABySS can be run across a range of k-mer lengths, and assemblies can be combined to resemble multiple k-mer assemblies; however, we abstained from that approach as it would have required sample-specific adjustments, which were not feasible within the scope of our study. Running Trinity with default parameters failed for some samples and workflows, and error messages indicated inappropriate default RAM settings. Despite thorough efforts to run Trinity with adjusted RAM settings as recommended by the developers (https://trinityrnaseq.github.io/performance/mem.html), we ultimately failed to automatize Trinity to run consistently across all samples and workflows. Manual adjustments for each sample and workflow might have resulted in successful Trinity assemblies; however, these were also not feasible within the scope of our study. Therefore, we excluded Trinity from the analysis.

#### Step four (mapping)

We employed two programs to map reads to assembled scaffolds to determine the read abundance of each scaffold: BWA v0.7.17 (Li and Durbin, 2009) and Bowtie2 v2.3.3.1 (Langmead and Salzberg, 2012) using default parameters. We processed mapped reads using the coverage function of samtools v1.10 (Li et al., 2009) to obtain the mean per-base coverage of each scaffold.

#### Step five (taxonomic annotation)

We used the SILVA132_NR99 SSU and LSU reference databases for taxonomic annotations (Quast et al., 2013), downloaded on 28 Aug 2020. NCBI’s Genbank database is also often used for taxonomic annotations of metagenomics data; however, initial tests showed that results were substantially skewed towards metagenomics-based workflows because entire genomes of the mock community taxa are available on Genbank. In the context of environmental sampling, at this point, it is unlikely that reference databases contain the full genomes of all sampled taxa. Therefore, to allow for a more realistic comparison of metagenomics and total RNA-Seq in an environmental context, we only used the SILVA database.

While there are many categories of taxonomic annotation tools, we limited our benchmarking to tools based on sequence similarity methods, which require reference databases, and explored two common tools: kraken2 v2.1.1 (Wood et al., 2019), based on k-mer matching, and BLAST v2.10.0 (Altschul et al., 1990), based on local alignments. We applied BLAST in two different ways, therefore using three approaches in total to classify each scaffold taxonomically: 1) k-mer-based using kraken2 with default parameters; 2) alignment-based using BLAST without further filtering of taxonomy hits; and 3) alignment-based using BLAST with further filtering of taxonomy hits. BLAST was run with an E-value cut-off of e-05 and otherwise default parameters. When running BLAST without further filtering of taxonomy hits, we kept the taxonomy hit with the highest bitscore per sequence unless there were multiple taxonomy hits with an identical highest bitscore, in which case we kept the lowest common ancestor (LCA). To filter BLAST taxonomy hits, we followed steps applied by the programs CREST (Lanzén et al., 2012) and BASTA (Kahlke and Ralph, 2019): filtering out taxonomy hits below a bitscore of 155 and an alignment length of 100, only keeping taxonomy hits within 2% of the best bitscore of each sequence, applying a cut-off for taxonomic ranks based on BLAST percent identity values (species: 99%, genus: 97%, family: 95%, order: 90%, class: 85%, phylum: 80%), and identifying the LCA. The setup of BLAST and kraken2 databases for SILVA required manual adaptations, which are described in Supplemental Material 2.

### Statistical analysis

#### Preprocessing

For the statistical analysis, all workflow results were further processed in Python v3.7.9 (Van Rossum and Drake, 2009). The full Python code is available on GitHub (https://github.com/hempelc/metagenomics-vs-totalRNASeq) and uses the modules Pandas v1.0.5 (Reback et al., 2020) and NumPy v1.19.1 (Harris et al., 2020). The evaluation was carried out at genus and species rank, respectively, to evaluate the accuracy and significance of workflows and tools on different taxonomic resolutions. Higher ranks were excluded from the evaluation since multiple species in the mock community were from the same family and therefore not distinguishable at higher ranks. We determined the per-base coverage of each detected taxon for each workflow as follows: we selected all scaffolds assigned to each taxon respectively, multiplied their mean per-base coverage by their length to determine their total number of covered bases, summarized the length and the total number of covered bases across all scaffolds, respectively, and divided the summarized total number of covered bases by the summarized length. That way, the per-base coverage of taxa reflected their cumulative scaffold length. To account for cross-contamination during filtration and extraction, we ran all workflows on the negative controls that were co-filtered and co-extracted with the mock community and subtracted the resulting taxa per base coverages of both controls from the same workflows utilized for the mock community. We then converted per base coverages into relative abundances by normalizing them so that they added up to 1.

#### Accuracy determination of workflows

Accuracy was determined independently for the three replicates. To determine the accuracy of each workflow within each replicate, we first generated accuracy metrics for each workflow. We considered the relative abundance of each expected taxon, i.e., each taxon in the mock community, as well as the relative abundance of each false-positive taxon introduced across all workflows and classifications as *NA*, i.e., no possible classification. Furthermore, we converted relative abundances into presence-absence (P-A) data to determine the accuracy based on both data types and generated separate accuracy metrics for relative abundances and P-A data.

We generated abundance-based accuracy metrics as follows: we used the observed relative abundance of each expected taxon as an individual accuracy metric and the summed observed relative abundance of all false-positive taxa as an additional metric. We defined P-A data-based accuracy metrics as follows: *true positive* (TP), number of taxa that were expected and observed, and *false positive* (FP), number of taxa that were not expected but observed. The latter included classifications as *NA*, i.e., no possible classification. We did not define *true negative* and *false negative* metrics since in our specific case they only provided redundant information for accuracy determination based on Euclidean distances, as described below.

Then, we converted the expected abundances and expected P-A of the mock community taxa and false-positive taxa, including classifications as *NA*, into accuracy metrics to resemble an unbiased workflow, which served as a reference to determine the accuracy of each workflow. For the mock community taxa, we used relative abundances based on genome copy number as given by the manufacturer for expected relative abundances (Table 1). The average relative abundance deviation of mock community taxa was < 30% according to the manufacturer. For false-positive taxa and *NA*, we set the expected relative abundances to zero before generating the expected metric.

Given that the abundance-based metrics were compositional, we followed appropriate steps for analyzing compositional data as pointed out by Gloor et al. (2017). Therefore, we applied simple multiplicative replacement to replace zeros among abundance-based metrics across all workflows, including the reference, using the multiplicative_replacement function of the python module scikit-bio v0.5.6 (scikit-bio development team, 2020). The function replaces zeros with a small positive value δ, which is based on the number of components, i.e., metrics while ensuring that compositions still add up to 1. Since the default δ was higher than the lowest expected abundance and therefore inappropriate, we manually defined δ as three orders of magnitude below the lowest expected abundance. Then, we applied a centred log-ratio (clr) transformation using the clr function of scikit-bio, which captures the relationships between features, i.e., metrics and makes the data symmetric and linearly related.

The accuracy of each workflow was determined by calculating the Euclidean distance between the metrics of each workflow and the expected reference metrics. The smaller the distance was, the more similar was the workflow to the reference, and the higher was the accuracy. The Euclidean distance of clr-transformed data is defined as the Aitchison distance (Aitchison et al., 2000).

#### Determining the most accurate data-processing tools for total RNA-Seq and metagenomics

We determined the most accurate tools for both total RNA-Seq and metagenomics by selecting the tool with the smallest minimum distance to the reference, representing the highest similarity to the mock community, for each workflow and each sequence processing step (Fig. 2)

To further evaluate the most similar workflows to the reference, we clustered workflows based on their distance to the reference. Since we clustered based on only one feature, distance, we applied a gaussian kernel density estimation using the KernelDensity function of scikit-learn to identify local minima of the distance distribution and split workflows into intervals, i.e., clusters based on local minima. We used a grid search to identify the most appropriate value for the bandwidth parameter using the GridSearchCV function of scikit-learn. We identified the closest cluster to the reference and determined the relative frequency of each tool among all workflows in the closest cluster for each processing step.

Lastly, to identify if tools overall had a significant impact on accuracy, we evaluated if they were significantly related to the distance to the reference. We converted tools into binary dummy variables and tested for significant correlations between each tool and the distance to the reference by calculating the point-biserial correlation coefficient, which measures the strength of the association between continuous and binary variables, using the pointbiserialr function of the python module SciPy v1.7.1 (Virtanen et al., 2020).

#### Comparing the accuracy of metagenomics and total RNA-Seq at equal sequencing depth

To test whether total RNA-Seq provided more accurate taxonomic identification of microbial communities than metagenomics at equal sequencing depths, we compared the accuracy metrics and Euclidean distance between the most accurate workflows of the three replicates. Since metagenomics replicates showed a much higher read depth than total RNA-Seq replicates, we subsampled metagenomics replicates to equal sequencing depths. DNA and RNA were co-extracted from the water samples, so we subsampled each metagenomics replicate to the read depth of the corresponding total RNA-Seq replicate. Random subsampling was performed ten times, and the most accurate workflow for each metagenomics replicate identified in our previous accuracy analysis was re-run for each subsample. When multiple workflows generated identical results, we re-ran the workflow with the lowest runtime. We generated accuracy metrics as described above and determined the mean average of each metric among the ten subsamples for each replicate. For P-A data, we rounded the averaged metrics to integers to avoid fractional numbers among the TP and FP metrics. Then, we calculated the Euclidean distance of the averaged accuracy metrics to the reference metrics as described above. We compared the accuracy metrics and Euclidean distance of the three subsampled metagenomics replicates to those of the most accurate workflows of the total RNA-Seq replicates. Lastly, we tested for significant differences between metagenomics- and total RNA-Seq-based Euclidean distances. Since DNA and RNA were co-extracted from the same samples and therefore not independent, we applied a paired t-test between the Euclidean distances of the three metagenomics and total RNA-Seq replicates using the ttest_rel function of SciPy.

#### Evaluating the relationship between sequencing depth and accuracy

To test if at increasing sequencing depths, the accuracy of total RNA-Seq increased faster than that of metagenomics, we evaluated the relationship between sequencing depth and accuracy by subsampling replicates at read depths of 1,000, 2,500, 5,000, 10,000, 20,000, 40,000, 60,000, 78,149, 94,633, 120,144, 200,000, 300,000, 400,000, 500,000, 600,000, 644,634, 669,382, and 817,619 reads and running the most accurate workflow for each replicate based on the highest similarity to the mock community. The maximum observed number of reads among all replicates exceeded the available number of reads for some replicates, and therefore, subsampling was only done up to the total number of reads in each replicate. We calculated the Euclidean distance to the reference for each replicate at each subsampled read depth and calculated the mean Euclidean distance across the subsamples. Euclidean distances were comparable between metagenomics and total RNA-Seq up to a read depth of 94,633, 78,149, and 120,144 reads for the first, second, and third replicate, respectively, and we generated ordinary least squares regression lines for each replicate of both approaches based on the mean Euclidean distances of the comparable data using the OLS function of the python module statsmodels v0.12.1 (Seabold and Perktold, 2010). Results based on abundance data gave impractical results for the generation of regression lines below 40,000 reads. At such low sequencing depths, often no single taxon was found, and even if some were found, the random subsampling was objected to high amounts of variance, which strongly skewed mean Euclidean distances based on abundance data, causing them to initially go up rather than down in some replicates. Therefore, we excluded these small subsamples when generating regression lines for abundance-based evaluations. For visualization purposes, we further generated the overall mean Euclidean distance and standard deviation (SD) across the subsamples of all metagenomics and total RNA-Seq replicates, respectively, and generated ordinary least squares regression lines based on the overall mean Euclidean distance. To test if the accuracy of total RNA-Seq increased significantly faster than that of metagenomics at increasing sequencing depths, we applied a paired t-test between the regression curve coefficients of the three metagenomics and total RNA-Seq replicates using the ttest_rel function of SciPy. To test if total RNA-Seq overall significantly improved the accuracy in comparison to metagenomics, we generated two linear models, one nested inside the other. One model involved the read depth as an independent variable to predict the Euclidean distance as the dependent variable, and the other model additionally involved the sequencing method (metagenomics/total RNA-Seq) as a binary independent variable:

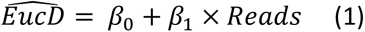

and

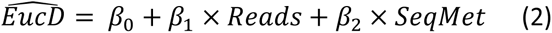

where *EucD* is the Euclidean distance, *Reads* is the read depth, and *SeqMet* is the sequencing method. By applying a partial *F*-test, we tested if the addition of the sequencing method as a binary independent variable significantly improved the Euclidean distance prediction, which indicated if one sequencing method was significantly more accurate than the other. We manually generated the models using the ols function of statsmodel and generated the *F*-statistic by performing an ANOVA for linear models using the anova_lm function of statsmodel.

## RESULTS

### Sequencing results

We obtained 2,428,038 paired-end reads for the mock community samples and controls (Bioproject number: PRJNA819997, SRA accession number: SRR18488964–SRR18488973). Since we normalized DNA and RNA samples based on volume during library preparation, which included samples from a different project with higher DNA/RNA concentrations, the number of reads that samples among DNA and RNA libraries received was dependent on the concentrations of the other normalized samples. The ratio between mock community samples and other samples seemed to substantially vary between DNA and RNA samples, and DNA samples received many more reads than RNA samples (710,545 reads on average for DNA samples vs. 97,642 reads on average for RNA samples (Fig. S1)). Therefore, DNA samples were randomly subsampled to allow comparisons at comparable read depths, as explained in the Methods section “Subsampling”. Notably, most of the RNA reads consisted of duplicates, potentially due to identical ribosomal sequences, whereas duplicates made up only a small portion of DNA reads. All negative controls contained a few reads (Fig. S1) and were processed in the same way as the samples. Negative control reads were subtracted from the samples to remove cross-contamination, although this did not impact the presented results.

### Most accurate data-processing tools for total RNA-Seq and metagenomics

Results differed substantially between replicates and evaluation levels, i.e., the taxonomic rank (species/genus) and data type (abundance/P-A) used for the evaluation (Fig. 3). Each column in Figure 3 represents one evaluation level utilized for one of three replicates and shows the relative proportions of data-processing tools within the cluster of workflows with the highest mean accuracy, the significance of correlations between tools and accuracy, and the most accurate tools. The frequency of tools and the most accurate tool were determined for each processing step separately.

**Figure 3:**
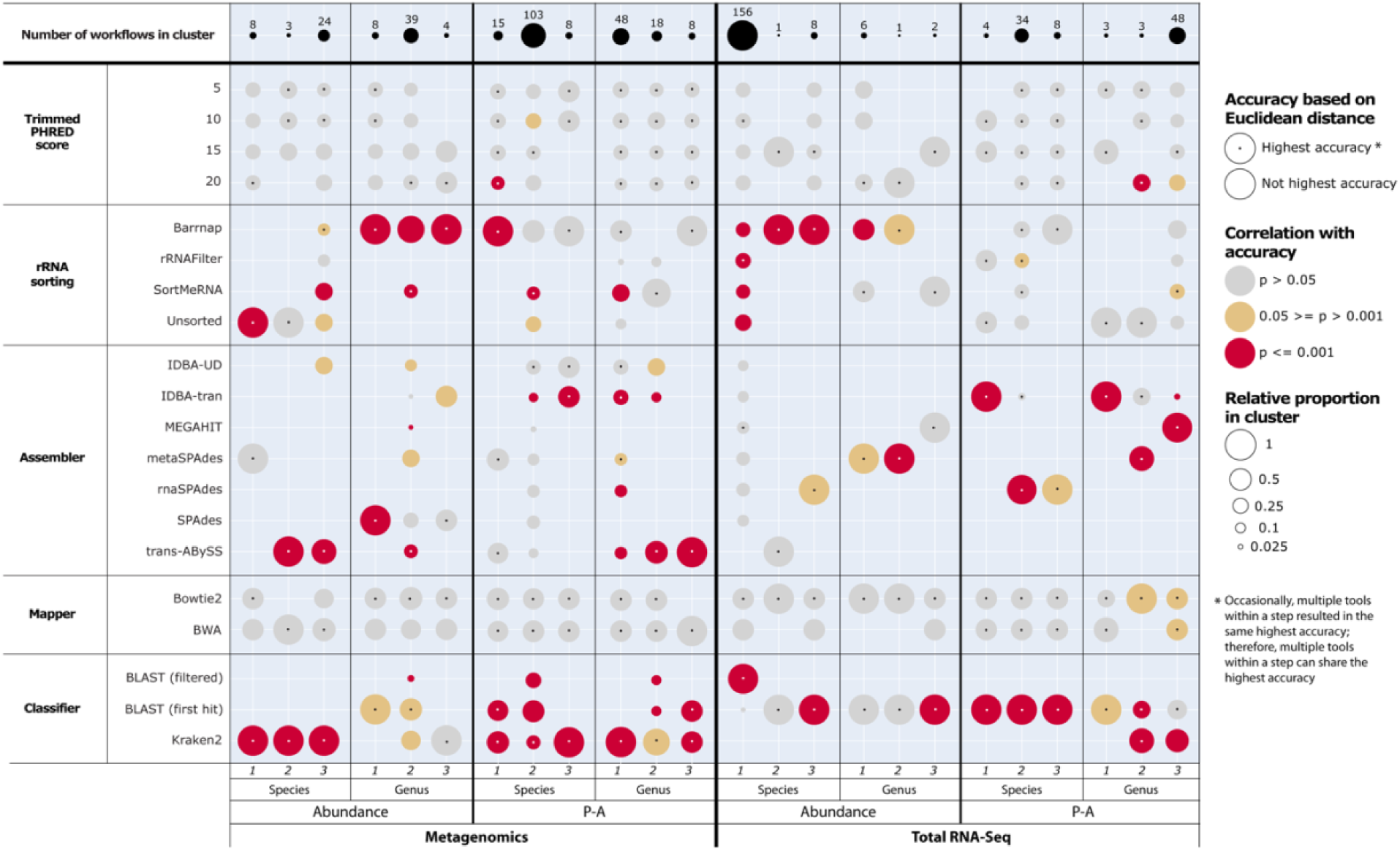
Relative frequency of data-processing tools within clusters of most accurate workflows (circle size), significance of correlations between tools and accuracy (circle colour), and most accurate tools based on different evaluation levels (dot in circle centre). Evaluation levels consisted of combinations of sequencing type (metagenomics/total RNA-Seq), data type (abundance/P-A), and taxonomic rank (genus/species). Each column represents one evaluation level utilized for one of three replicates. The relative frequency of tools and the tool with the highest accuracy were determined for each data-processing step separately. Performances differed substantially among replicates and evaluation levels.

The relative frequency of tools within the most accurate clusters denoted dominance of specific tools in the clusters, indicated by a high frequency of a single tool. The relative frequencies of utilized PHRED scores and mapping tools were overall evenly distributed, indicating that the most accurate workflows were independent of the utilized PHRED score and mapping tools, except for some small clusters. For all other processing steps, clusters were dominated by one or two tools in most cases. However, the dominating tools varied substantially among replicates, with a few exceptions, notably Kraken2 dominating the most accurate cluster of metagenomics-based workflows for species-abundance evaluations, Barrnap dominating the most accurate cluster of metagenomics-based workflows for genus-abundance and species-P-A evaluations, and BLAST (first hit) dominating the most accurate cluster of total RNA-Seq-based workflows for genus-abundance and species-P-A evaluations. No preferable tools across all evaluation levels could be identified for the assembly step, and there was no relationship between metagenomics-based workflows and metagenomics-optimized assemblers or total RNA-Seq-based workflows and or total RNA-Seq-optimized assemblers. BLAST (filtered) performed poorly across all evaluation levels except for a few replicates.

Significance indicates whether tools correlate with consistently higher accuracy across all utilized workflows and, therefore, consistently perform better. Non-significant correlations (p > 0.05) did not indicate poor performance but rather that the performance was also dependent on the other tools used within workflows. Utilized PHRED scores and mapping tools showed no significant correlation to accuracy, with only a few exceptions. Notable patterns were that Kraken2 overall significantly correlated with higher accuracies in metagenomics-based workflows and that BLAST (first hit) overall significantly correlated with higher accuracy in total RNA-Seq-based workflows.

The single most accurate tool varied among replicates and evaluation levels but was often the tool that dominated the cluster. Notably, for P-A-based evaluations, often multiple tools within a step performed identically. This was most notable for utilized PHRED scores and mapping tools, further confirming that accuracy was independent of the utilized tools within these two steps. Kraken2 was among the tools with the highest accuracy in almost all metagenomics-based workflows, and the same was true for BLAST (first hit) for total RNA-Seq-based workflows.

Overall, the most accurate tools and workflows depended on evaluation levels and strongly varied within evaluation levels, and performances differed among replicates, indicating that the accuracy and significance of tools can vary from sample to sample, even across replicates of a highly controlled mock community. Kraken2 and BLAST (first hit) were mostly preferable for taxonomic classification within metagenomics and total RNA-Seq workflows, respectively. Utilized PHRED scores and mapping tools were interchangeable in terms of accuracy. Assemblers and rRNA sorting tools were extremely evaluation level- and replicate-dependent, however, no sorting or Barrnap were overall preferable over rRNAFilter and SortMeRNA for rRNA sorting.

### Comparing the accuracy of metagenomics and total RNA-Seq at equal sequencing depths

All replicates showed variations across all evaluation levels (Fig. 4). However, based on Euclidean distances to the reference, total RNA-Seq-based workflows were significantly more similar to the reference than metagenomics-based workflows for all evaluation levels (genus- abundance: p = 0.011, genus-P-A: p = 0.014, species-abundance: p = 0.019, species-P-A: p = 0.003), indicating that total RNA-Seq-based workflows outperformed metagenomics-based workflows in terms of accuracy if appropriate data-processing tools were used.

**Figure 4:**
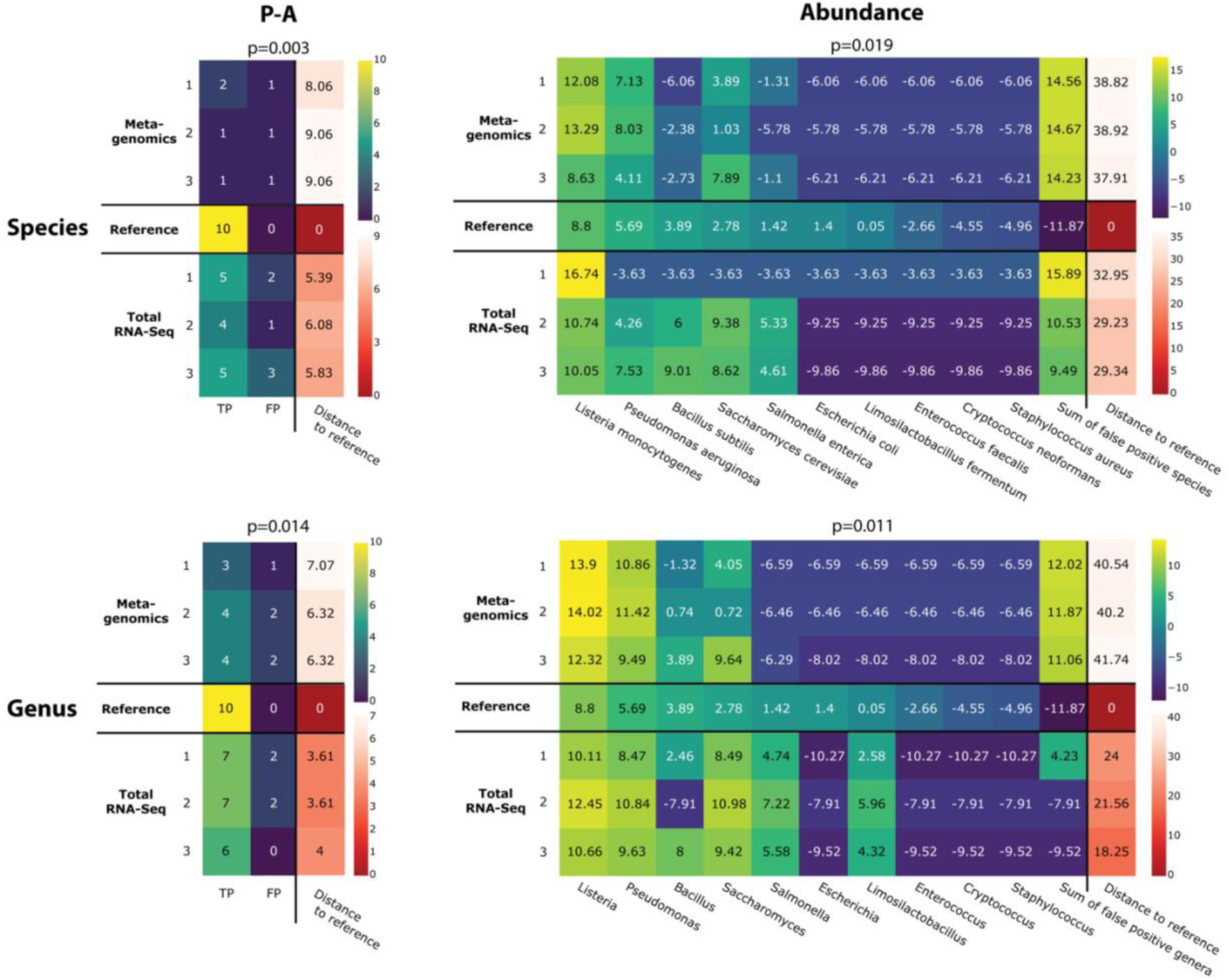
Comparison of the most accurate metagenomics- and total RNA-Seq-based workflow of each replicate for multiple evaluation levels based on the Euclidean distance to the reference. Evaluation levels consisted of combinations of data type (abundance/P-A), and taxonomic rank (genus/species). Since individual metrics were on a different scale than Euclidean distance to the reference, two different colour scales were applied. The reference in the middle row of the heatmaps represents expected metrics, and the closer metagenomics or total RNA-Seq metrics were to the reference, the more accurate they were. Abundance-based metrics underwent multiplicative replacement followed by clr-transformation. p-values are based on paired t-tests between metagenomics- and total RNA-Seq-based Euclidean distances to the reference. All replicates showed variations across all evaluation levels; however, total RNA-Seq-based workflows were significantly more similar to the reference than metagenomics-based workflows (p < 0.05) for all evaluation levels. For P-A-based evaluations, absolute differences among metrics and replicates were small (left). Metagenomics- and total RNA-Seq-based replicates failed to detect the 5 or 6 taxa with the lowest abundance (right).

For P-A-based evaluations, absolute differences among metrics and replicates were small (Fig. 4, left). Replicate Total RNA-Seq 1 was most accurate, detecting 5 and 7 out of 10 expected genera/species (TP = 5 and 7) and detecting only 2 false positive genera and species, respectively (FP = 2).

When comparing individual metrics within abundance-based evaluations, all metagenomics-based replicates introduced a substantial abundance of false-positive genera, while this was only the case for one in three total RNA-Seq-based replicates (Fig. 4, bottom right). In contrast, all metagenomics- and total RNA-Seq-based replicates introduced a substantial abundance of false-positive species (Fig. 4, top right). Further analysis of the abundance and composition of false-positive species revealed that they were mostly made up of *NA*, i.e., no classification (on average 23.7% and 22.7% of all metagenomics and total RNA-Seq reads, respectively), while the contribution of individual false-positive species was comparably small (on average 0.13–1.6% and 0.01–7.9% of all metagenomics and total RNA-Seq reads, respectively). However, given the extremely low expected abundance of most expected taxa, these results showed that the abundance of false-positive species was still considerably higher than that of most expected taxa.

Abundance-based metrics spanned five orders of magnitude based on genome copy number as given by the manufacturer (Tab. 1), and all metagenomics- and total RNA-Seq-based replicates failed to detect the 4–6 taxa with the lowest abundance. Notably, total RNA-Seq detected 2 genera more than metagenomics, of which one had an abundance one order of magnitude lower than that of a non-detected genus. This indicated that abundance was not the only factor for detection and that other mechanisms affected if a genus was detected.

The most accurate workflow for replicate Total RNA-Seq 1 based on species-abundance metrics only detected the most abundant species, *Listeria monocytogenes*, and no other taxon, indicating that the accuracy of all other workflows utilized for this replicate was lower and, therefore, that not detecting 9/10 species was more accurate in terms of Euclidean distance to the reference than detecting multiple expected taxa with biased abundances, as was the case for the other workflows with lower accuracy.

### Relationship between sequencing depth and accuracy

The relationship between sequencing depth and accuracy varied among evaluation levels and sequencing types (Fig. 5). Notably, for abundance-based evaluations, the mean Euclidean distance to the reference increased for subsamples up to 20,000 reads. This observation was due to more expected taxa being found at increasing sequencing depth, but initially at abundances that were more biased than when expected taxa were not found at all, which initially decreased the accuracy. This made subsamples up to 20,000 reads impractical for the regression curve- based comparison of total RNA-Seq and metagenomics, and hence we excluded them, but the respective curves are shown in Figure S2.

**Figure 5:**
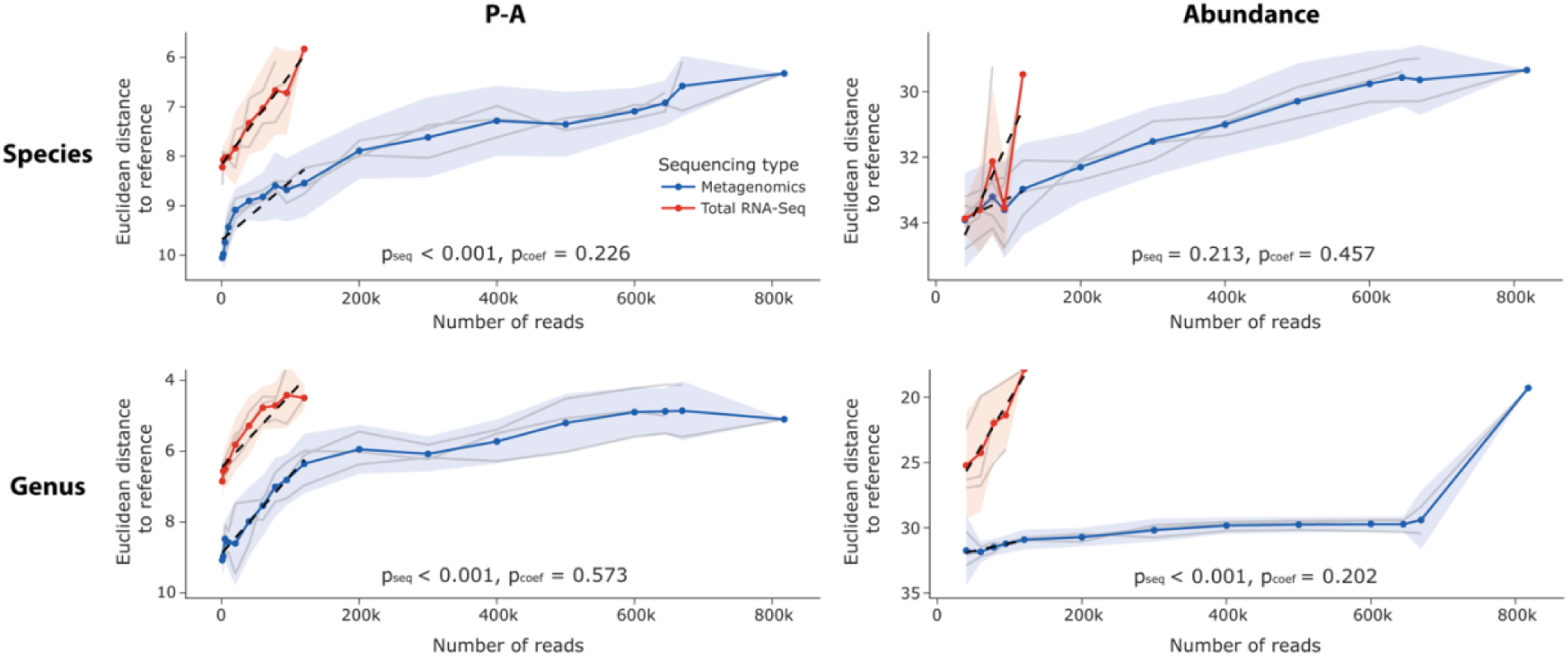
Relationship between sequencing depth and accuracy for multiple evaluation levels. Evaluation levels consisted of combinations of data type (abundance/P-A), and taxonomic rank (genus/species). Blue and red lines indicate the mean Euclidean distance of all metagenomics and total RNA-Seq replicates, which have each been subsampled ten times, and the area around the lines indicates the standard deviation (SD). Lower Euclidean distances are a proxy for higher accuracy. The y-axis is inverted, and its scale varies among graphs. The SD equals zero at the highest number of reads since all available reads were used and, therefore, no subsamples could be generated. Individual replicates are shown as grey lines. Regression curves are shown as dashed black lines for the portion of the data that was comparable between metagenomics and total RNA-Seq. p_seq_-values are based on partial F-tests between linear models including or excluding the sequencing method (metagenomics/total RNA-Seq) as a binary independent variable. p_coef_-values are based on paired t-tests between the coefficients of regression curves of individual metagenomics and total RNA-Seq replicates based on the comparable portion of the data.

For all evaluation levels but genus-P-A, the mean accuracy of total RNA-Seq increased faster than that of metagenomics. However, when testing for significantly faster increases in accuracy of total RNA-Seq replicates in comparison to metagenomics replicates, i.e., significant differences in the coefficients of the regression curves, no significant differences were found (genus-abundance: p_coef_ = 0.202, genus-P-A: p_coef_ = 0.573, species-abundance: p_coef_ = 0.457, species-P-A: p_coef_ = 0.226). This might be explained by the high variations among replicates (Fig. 5, grey lines). While no statistically significant differences could be confirmed, the observed trend of the regression curves indicates that with more replicates, statistically significant differences might become apparent.

Nevertheless, the mean accuracy of total RNA-Seq was consistently higher than that of metagenomics for all evaluation levels but species-abundance, even at sequencing depths approximately one order of magnitude lower than that of metagenomics. For the evaluation level species-abundance, the mean accuracy of total RNA-Seq was initially lower than that of metagenomics but increased rapidly until they were approximately equal. Partial *F*-tests between linear models including or excluding the sequencing method (metagenomics/total RNA-Seq) as a binary independent variable confirmed that for all evaluation levels but species-abundance, the addition of the sequencing method into the models significantly improved Euclidean distance prediction (genus-abundance: p_seq_ < 0.001, genus-P-A: p_seq_ < 0.001, species-abundance: p_seq_ = 0.213, species-P-A: p_seq_ < 0.001). These results confirmed that total RNA-Seq was significantly more accurate than metagenomics.

## DISCUSSION

Our first aim was to test which data-processing workflow was the most accurate for total RNA-Seq and if it coincided with the most accurate data-processing workflow for metagenomics. We tested 736 different workflows since many studies highlight that HTS-based results are heavily influenced by the tools used to process the data (Bashiardes et al., 2016; Knight et al., 2018; McIntyre et al., 2017; Quince et al., 2017; Shakya et al., 2019; Vollmers et al., 2017), and we assumed that different tools would perform most accurately for metagenomics and total RNA-Seq data, respectively, since both sequencing methods result in different read compositions.

Our results showed that for the steps quality filtering and trimming, rRNA sorting, and mapping, the most accurate tools were similar for metagenomics and total RNA-Seq, and the choice of quality filtering, trimming, and mapping tools had overall no impact on the accuracy for both sequencing methods. Only for the classification step, the most accurate tools differed, with Kraken2 being overall the most accurate for metagenomics and BLAST (first hit) for total RNA-Seq. However, differences among evaluation levels and replicates were apparent, especially for assemblers, indicating that the most accurate tools were dependent on evaluation level and replicate rather than sequencing type. These results show that certain processing steps need more attention than others - in the context of our study, this would refer to the utilized rRNA sorting tools, assemblers, and classifiers, while choices for trimming, quality filtering, and mapping might require less attention. It should be noted, though, that the quality of our sequences was almost exclusively over PHRED 30 according to mean and per sequence quality scores (Fig. S3), so trimming and quality filtering might not have had a big effect in our study but could still have significant effects in studies yielding lower-quality data. Furthermore, the fact that the most accurate tools were highly dependent on the replicates indicated that the communities varied among replicates. The microbial mock community used in our study has an average relative abundance deviation of maximal 30% according to the manufacturer, meaning that deviations from the theoretical composition are possible. Given that the majority of the mock community taxa had an extremely low abundance, and that additional bias was likely introduced when splitting the mock community into three replicates as well as during filtration, extraction, and sequencing, it is not surprising that results differed somewhat among replicates. Further mock community-based tests involving different taxa, more replicates, and ideally, less variability among replicates are required to compare the most appropriate data-processing tools for metagenomics and total RNA-Seq.

To our knowledge, the scale of our benchmarking approach is unique in the scope of tested combinations of diverse data-processing tools. However, while it allowed us to analyze the impact of individual data-processing steps and tools on microbial identification accuracy, it did not include tests on multiple datasets with different taxonomic compositions. Furthermore, since we only tested a selection of best-supported and most commonly applied tools from prior studies, the number of tools included for each step was low in comparison to benchmark studies that focus on individual processing steps (Hleap et al., 2021; Hölzer and Marz, 2019; McIntyre et al., 2017; Sierra et al., 2020; Vollmers et al., 2017). This also permits testing parameter modifications of specific tools; e.g., Hleap et al. (2021) applied a grid search approach to test multiple parameters of each tool and selected the most appropriate parameters, while Vollmers et al. (2017) tested multiple assemblers with two different k-mer lengths. Such fine-tuning was not feasible within the scope of our study but might increase the performance of specific tools. Nevertheless, for the identification of the impact of specific processing steps and to compare the most accurate data-processing tools for metagenomics and total RNA-Seq, our benchmarking study provides important information on a broader scale.

We detected no difference in accuracy for metagenomics-optimized assemblers using metagenomics data or for metatranscriptomics-optimized assemblers using total RNA-Seq data. Metatranscriptomics-optimized assemblers were developed to overcome the issue of uneven expression levels among RNA-Seq data, which hampers the assembly of lowly expressed transcriptomes. Although many of these assemblers have been developed and tested on RNA-Seq data, only once were they reciprocally tested using metagenomics data. Bushmanova et al. (2019) tested their metagenomics-optimized SPAdes assembler with RNA-Seq data, with similar performance but a tendency to generate longer contigs incorrectly and an inability to detect isoforms of mRNAs. Based on this and our own findings, we suggest using established assemblers that have been validated in metagenomics-specific benchmarking studies, such as metaSPAdes and MEGAHIT (Awad et al., 2017; Quince et al., 2017; van der Walt et al., 2017).

Based on our central idea that total RNA-Seq recovers more taxonomically informative sequences than metagenomics, our second aim was to test if total RNA-Seq provides more accurate taxonomic identifications than metagenomics at equal sequencing depth. This was corroborated by our results, raising the potential to increase the accuracy of taxonomic identifications of diverse microbial communities when applying targeted-PCR-free shotgun sequencing. This, in turn, could increase the effectiveness of freshwater assessments, since microbes respond faster to environmental changes and, therefore, might better represent environmental conditions than other taxa, as suggested for prokaryotes (McArthur, 2001; Smith et al., 2015), unicellular eukaryotes (Foissner and Berger, 1996; Pawlowski et al., 2016; Payne, 2013; Stoeck et al., 2018), or both (Cordier et al., 2019; Sagova-Mareckova et al., 2021).

Our third aim was to test if the accuracy of total RNA-Seq increased faster than that of metagenomics with increasing sequencing depth due to greater taxonomic signal in the recovered total RNA-Seq reads. While this was not supported by our results, the accuracy of total RNA-Seq was comparable to or even outperformed that of metagenomics at sequencing depths almost one order of magnitude lower. Furthermore, our results indicate that if we had increased the number of replicates, our assumption might have been confirmed. Leese et al. (2018) noted that cost-effectiveness needs to be considered when proposing new methods for freshwater assessment and that targeted-PCR-free techniques lack sufficient validation and proper reference data for routine applications despite their huge potential. Our study shows that total RNA-Seq might represent the sought-after, cost-efficient, and targeted-PCR-free method worthy of further exploration as it can be effective at substantially lower sequencing depth and, therefore, substantially reduce costs. Traditional metatranscriptomics is also gaining increasing attention for freshwater assessments. It has been suggested that microbial function might be a better proxy for environmental change than taxon identity because functional shifts can occur before taxonomic shifts (Cordier et al. 2019, 2020a). This growing interest in metatranscriptomics could also increase the applicability of total RNA-Seq as both approaches could be performed complementarily using the same RNA extracts. This would have the added benefit of being able to study taxon-function relationships. However, there have also been concerns regarding the applicability of incorporating RNA into routine assessments due to the instability of RNA and higher costs of RNA sample collections (Cordier et al., 2019). But recent studies of environmental RNA (eRNA) suggest that it is much more stable in the environment than previously assumed, making it indeed suitable for routine assessments and opening possibilities for an entirely new field of environmental research, called environmental transcriptomics (Cristescu, 2019; Yates et al., 2021).

## Conclusion

Our study demonstrates that the impact of data-processing tools on metagenomics and total RNA-Seq data substantially varies among replicates and evaluation levels. Furthermore, metagenomics-optimized assemblers do not uniformly improve metagenomics data and neither do metatranscriptomics-optimized assemblers for total RNA-Seq data. Further studies with a higher resolution of specific data-processing steps are required to finetune the most appropriate choices for a given context.

Total RNA-Seq provided more accurate taxonomic identifications for our microbial mock community than metagenomics at equal sequencing depths and even at sequencing depths one order of magnitude lower. These results indicate that total RNA-Seq represents a good alternative to metagenomics when it comes to targeted-PCR-free taxonomic identifications of microbial communities and that a substantial reduction in sequencing costs might be possible while maintaining accuracy. This could benefit routine freshwater assessments, which require cost-effective methods, and allows for the incorporation of microbes into freshwater assessments. In the context of current eRNA and environmental transcriptomics research, total RNA-Seq could be a complementary approach to metatranscriptomics, allowing the establishment of taxon-function relationships. Further research on environmental samples is required to confirm the advantages of total RNA-Seq over metagenomics in applied settings.

## DATA AVAILABILITY

The sequencing data is available under Bioproject number PRJNA819997 and SRA accession numbers SRR18488964–SRR18488973.

## ACKNOWLEDGEMENTS

We would like to thank Karl Cottenie, Anders Lanzen, Florian Leese, and Nicole Ricker for helpful advice and stimulating discussions.

## REFERENCES

Aitchison, J., Barceló-Vidal, C., Martín-Fernández, J. A., and Pawlowsky-Glahn, V. (2000). Logratio analysis and compositional distance. Math. Geol. 32, 271–275. doi:10.1023/A:1007529726302.

Alberdi, A., Aizpurua, O., Gilbert, M. T. P., and Bohmann, K. (2018). Scrutinizing key steps for reliable metabarcoding of environmental samples. Methods Ecol. Evol. 9, 134–147. doi:10.1111/2041-210X.12849.

Almeida, O. G. G., and De Martinis, E. C. P. (2019). Bioinformatics tools to assess metagenomic data for applied microbiology. Appl. Microbiol. Biotechnol. 103, 69–82. doi:10.1007/s00253-018-9464-9.

Altschul, stephen F., Gish, W., Miller, W., Myers, E. W., and Lipman, D. J. (1990). Basic Local Alignment Search Tool. J. Mol. Biol. 215, 403–410.

Awad, S., Irber, L., and Brown, C. T. (2017). Evaluating Metagenome Assembly on a Simple Defined Community with Many Strain Variants. bioRxiv, 155358. doi:10.1101/155358.

Bang-Andreasen, T., Anwar, M. Z., Lanzén, A., Kjøller, R., Rønn, R., Ekelund, F., et al. (2020). Total RNA sequencing reveals multilevel microbial community changes and functional responses to wood ash application in agricultural and forest soil. FEMS Microbiol. Ecol. 96, 1–13. doi:10.1093/femsec/fiaa016.

Bankevich, A., Nurk, S., Antipov, D., Gurevich, A. A., Dvorkin, M., Kulikov, A. S., et al. (2012). SPAdes: A New Genome Assembly Algorithm and Its Applications to Single-Cell Sequencing. J. Comput. Biol. 19, 455–477. doi:10.1089/cmb.2012.0021.

Bashiardes, S., Zilberman-Schapira, G., and Elinav, E. (2016). Use of metatranscriptomics in microbiome research. Bioinform. Biol. Insights 10, 19–25. doi:10.4137/BBI.S34610.

Bellinger, E. G., and Sigee, D. C. (2015). Freshwater Algae - Identification, Enumeration and Use as Bioindicators. 2nd ed. Chichester, West Sussex: John Wiley & Sons Ltd.

Bolger, A. M., Lohse, M., and Usadel, B. (2014). Trimmomatic: A flexible trimmer for Illumina sequence data. Bioinformatics 30, 2114–2120. doi:10.1093/bioinformatics/btu170.

Burger, J. (2006). Bioindicators: A review of their use in the environmental literature 1970– 2005. Environ. Bioindic. 1, 136–144. doi:10.1080/15555270600701540.

Bushmanova, E., Antipov, D., Lapidus, A., and Prjibelski, A. D. (2019). rnaSPAdes: A de novo transcriptome assembler and its application to RNA-Seq data. Gigascience 8, 1–13. doi:10.1093/gigascience/giz100.

Carini, P., Marsden, P. J., Leff, J. W., Morgan, E. E., Strickland, M. S., and Fierer, N. (2016). Relic DNA is abundant in soil and obscures estimates of soil microbial diversity. Nat. Microbiol. 2, 1–6. doi:10.1038/nmicrobiol.2016.242.

Clarridge, J. E. (2004). Impact of 16S rRNA gene sequence analysis for identification of bacteria on clinical microbiology and infectious diseases. Clin. Microbiol. Rev. 17, 840–862. doi:10.1128/CMR.17.4.840-862.2004.

Cordier, T., Alonso-Sáez, L., Apothéloz-Perret-Gentil, L., Aylagas, E., Bohan, D. A., Bouchez, A., et al. (2020a). Ecosystems monitoring powered by environmental genomics: A review of current strategies with an implementation roadmap. Mol. Ecol., 0–2. doi:10.1111/mec.15472.

Cordier, T., Lanzén, A., Apothéloz-Perret-Gentil, L., Stoeck, T., and Pawlowski, J. (2019). Embracing Environmental Genomics and Machine Learning for Routine Biomonitoring. Trends Microbiol. 27, 387–397. doi:10.1016/j.tim.2018.10.012.

Cordier, T., Sáez, L. A., Apotheloz-Perret-Gentil, L., Aylagas, E., Bohan, D. A., Bouchez, A., et al. (2020b). Ecosystems Monitoring Powered by Environmental Genomics: A Review of Current Strategies with An Implementation Roadmap. Preprints. doi:10.20944/PREPRINTS202001.0278.V1.

Cristescu, M. E. (2014). From barcoding single individuals to metabarcoding biological communities: Towards an integrative approach to the study of global biodiversity. Trends Ecol. Evol. 29, 566–571. doi:10.1016/j.tree.2014.08.001.

Cristescu, M. E. (2019). Can Environmental RNA Revolutionize Biodiversity Science? Trends Ecol. Evol. 34, 694–697. doi:10.1016/j.tree.2019.05.003.

Deiner, K., Bik, H. M., Mächler, E., Seymour, M., Lacoursière-Roussel, A., Altermatt, F., et al. (2017). Environmental DNA metabarcoding: Transforming how we survey animal and plant communities. Mol. Ecol. 26, 5872–5895. doi:10.1111/mec.14350.

Elekwachi, C. O., Wang, Z., Wu, X., Rabee, A., and Forster, R. J. (2017). Total rRNA-seq analysis gives insight into bacterial, fungal, protozoal and archaeal communities in the rumen using an optimized RNA isolation method. Front. Microbiol. 8, 1–14. doi:10.3389/fmicb.2017.01814.

Foissner, W., and Berger, H. (1996). A User-Friendly Guide to the Ciliates. Freshw. Biol. 35, 375– 482.

Geisen, S., Tveit, A. T., Clark, I. M., Richter, A., Svenning, M. M., Bonkowski, M., et al. (2015). Metatranscriptomic census of active protists in soils. ISME J. 9, 2178–2190. doi:10.1038/ismej.2015.30.

Gloor, G. B., Macklaim, J. M., Pawlowsky-Glahn, V., and Egozcue, J. J. (2017). Microbiome datasets are compositional: And this is not optional. Front. Microbiol. 8, 1–6. doi:10.3389/fmicb.2017.02224.

Gomez-Silvan, C., Leung, M. H. Y., Grue, K. A., Kaur, R., Tong, X., Lee, P. K. H., et al. (2018). A comparison of methods used to unveil the genetic and metabolic pool in the built environment. Microbiome 6, 1–16. doi:10.1186/s40168-018-0453-0.

Grabherr, M. G., Haas, B. J., Yassour, M., Levin, J. Z., Thompson, D. A., Amit, I., et al. (2013). Trinity: reconstructing a full-length transcriptome without a genome from RNA-Seq data. Nat. Biotechnol. 29, 644–652. doi:10.1038/nbt.1883.Trinity.

Harris, C. R., Millman, K. J., van der Walt, S. J., Gommers, R., Virtanen, P., Cournapeau, D., et al. (2020). Array programming with {NumPy}. Nature 585, 357–362. doi:10.1038/s41586-020-2649-2.

Haury, J., Peltre, M.-C., Trémolières, M., Barbe, J., Thiébaut, G., Bernez, I., et al. (2006). A new method to assess water trophy and organic pollution – the Macrophyte Biological Index for Rivers (IBMR): its application to different types of river and pollution. Hydrobiologia 570, 153–158. doi:10.1007/s10750-006-0175-3.

Hleap, J. S., Littlefair, J. E., Steinke, D., Hebert, P. D. N., and Cristescu, M. E. (2021). Assessment of current taxonomic assignment strategies for metabarcoding eukaryotes. Mol. Ecol. Resour. 21, 2190–2203.

Hölzer, M., and Marz, M. (2019). De novo transcriptome assembly: A comprehensive cross-species comparison of short-read RNA-Seq assemblers. Gigascience 8, 1–16. doi:10.1093/gigascience/giz039.

IPBES (2019). Summary for policymakers of the global assessment report on biodiversity and ecosystem services of the Intergovernmental Science-Policy Platform on Biodiversity and Ecosystem Services., eds. S. Díaz, J. Settele, E. S. Brondízio, H. T. Ngo, M. Guèze, J. Agard, et al. Bonn, Germany Available at: https://doi.org/10.5281/zenodo.3553579.

Janda, J. M., and Abbott, S. L. (2007). 16S rRNA gene sequencing for bacterial identification in the diagnostic laboratory: Pluses, perils, and pitfalls. J. Clin. Microbiol. 45, 2761–2764. doi:10.1128/JCM.01228-07.

Kahlke, T., and Ralph, P. J. (2019). BASTA – Taxonomic classification of sequences and sequence bins using last common ancestor estimations. Methods Ecol. Evol. 10, 100–103. doi:10.1111/2041-210X.13095.

Karr, J. R. (1981). Assessment of Biotic Integrity Using Fish Communities. Fisheries 6, 21–27. doi:10.1577/1548-8446(1981)006.

Knight, R., Navas, J., Quinn, R. A., Sanders, J. G., and Zhu, Q. (2018). Best practices for analysing microbiomes. Nat. Rev. Microbiol. doi:10.1038/s41579-018-0029-9.

Kopylova, E., Noé, L., and Touzet, H. (2012). SortMeRNA: Fast and accurate filtering of ribosomal RNAs in metatranscriptomic data. Bioinformatics 28, 3211–3217. doi:10.1093/bioinformatics/bts611.

Krehenwinkel, H., Wolf, M., Lim, J. Y., Rominger, A. J., Simison, W. B., and Gillespie, R. G. (2017). Estimating and mitigating amplification bias in qualitative and quantitative arthropod metabarcoding. Sci. Rep. 7, 1–12. doi:10.1038/s41598-017-17333-x.

Kubiszewski, I., Costanza, R., Anderson, S., and Sutton, P. (2017). The future value of ecosystem services: Global scenarios and national implications. Ecosyst. Serv. 26, 289–301. doi:10.1016/j.ecoser.2017.05.004.

Langmead, B., and Salzberg, S. L. (2012). Fast gapped-read alignment with Bowtie 2. Nat. Methods 9, 357–359. doi:10.1038/nmeth.1923.

Lanzén, A., Jørgensen, S. L., Bengtsson, M. M., Jonassen, I., Øvreås, L., and Urich, T. (2011). Exploring the composition and diversity of microbial communities at the Jan Mayen hydrothermal vent field using RNA and DNA. FEMS Microbiol. Ecol. 77, 577–589. doi:10.1111/j.1574-6941.2011.01138.x.

Lanzén, A., Jørgensen, S. L., Huson, D. H., Gorfer, M., Grindhaug, S. H., Jonassen, I., et al. (2012). CREST - Classification Resources for Environmental Sequence Tags. PLoS One 7. doi:10.1371/journal.pone.0049334.

Leese, F., Bouchez, A., Abarenkov, K., Altermatt, F., Borja, Á., Bruce, K., et al. (2018). Why We Need Sustainable Networks Bridging Countries, Disciplines, Cultures and Generations for Aquatic Biomonitoring 2.0: A Perspective Derived From the DNAqua-Net COST Action. Adv. Ecol. Res. 58, 63–99. doi:10.1016/bs.aecr.2018.01.001.

Li, D., Liu, C. M., Luo, R., Sadakane, K., and Lam, T. W. (2015). MEGAHIT: An ultra-fast single-node solution for large and complex metagenomics assembly via succinct de Bruijn graph. Bioinformatics 31, 1674–1676. doi:10.1093/bioinformatics/btv033.

Li, F., and Guan, L. L. (2017). Metatranscriptomic profiling reveals linkages between the active rumen microbiome and feed efficiency in beef cattle. Appl. Environ. Microbiol. 83, 1–16. doi:10.1128/AEM.00061-17.

Li, F., Henderson, G., Sun, X., Cox, F., Janssen, P. H., and Guan, L. L. (2016). Taxonomic assessment of rumen microbiota using total rna and targeted amplicon sequencing approaches. Front. Microbiol. 7. doi:10.3389/fmicb.2016.00987.

Li, H. Seqtk. Available at: https://github.com/lh3/seqtk.

Li, H., and Durbin, R. (2009). Fast and accurate short read alignment with Burrows-Wheeler transform. Bioinformatics 25, 1754–1760. doi:10.1093/bioinformatics/btp324.

Li, H., Handsaker, B., Wysoker, A., Fennell, T., Ruan, J., Homer, N., et al. (2009). The Sequence Alignment/Map format and SAMtools. Bioinformatics 25, 2078–2079. doi:10.1093/bioinformatics/btp352.

Logares, R., Sunagawa, S., Salazar, G., Cornejo-Castillo, F. M., Ferrera, I., Sarmento, H., et al. (2014). Metagenomic 16S rDNA Illumina tags are a powerful alternative to amplicon sequencing to explore diversity and structure of microbial communities. Environ. Microbiol. 16, 2659–2671. doi:10.1111/1462-2920.12250.

MacManes, M. D. (2014). On the optimal trimming of high-throughput mRNA sequence data. Front. Genet. 5, 1–7. doi:10.3389/fgene.2014.00013.

McArthur, J. V (2001). “Bacteria as Biomonitors,” in Bioassessment and Management of North American Freshwater Wetlands, eds. R. B. Rader, D. P. Batzer, and S. A. Wissinger (Chichester, West Sussex: John Wiley & Sons), 249–261. doi:https://doi.org/10.1002/aqc.509.

McIntyre, A. B. R., Ounit, R., Afshinnekoo, E., Prill, R. J., Hénaff, E., Alexander, N., et al. (2017). Comprehensive benchmarking and ensemble approaches for metagenomic classifiers. Genome Biol. 18, 1–19. doi:10.1186/s13059-017-1299-7.

Nurk, S., Meleshko, D., Korobeynikov, A., and Pevzner, P. A. (2017). MetaSPAdes: A new versatile metagenomic assembler. Genome Res. 27, 824–834. doi:10.1101/gr.213959.116.

Pawlowski, J., Audic, S., Adl, S., Bass, D., Belbahri, L., Berney, C., et al. (2012). CBOL Protist Working Group: Barcoding Eukaryotic Richness beyond the Animal, Plant, and Fungal Kingdoms. PLoS Biol. 10, e1001419. doi:10.1371/journal.pbio.1001419.

Pawlowski, J., Lejzerowicz, F., Apotheloz-Perret-Gentil, L., Visco, J. A., and Esling, P. (2016). Protist metabarcoding and environmental biomonitoring: Time for change. Eur. J. Protistol. 55, 12–25. doi:10.1016/j.ejop.2016.02.003.

Payne, R. J. (2013). Seven reasons why protists make useful bioindicators. Acta Protozool. 52, 105–113. doi:10.4467/16890027AP.13.0011.1108.

Peano, C., Pietrelli, A., Consolandi, C., Rossi, E., Petiti, L., Tagliabue, L., et al. (2013). An efficient rRNA removal method for RNA sequencing in GC-rich bacteria. Microb. Inform. Exp. 3, 1– 11. doi:10.1186/2042-5783-3-1.

Peng, Y., Leung, H. C. M., Yiu, S. M., and Chin, F. Y. L. (2012). IDBA-UD: A de novo assembler for single-cell and metagenomic sequencing data with highly uneven depth. Bioinformatics 28, 1420–1428. doi:10.1093/bioinformatics/bts174.

Peng, Y., Leung, H. C. M., Yiu, S. M., Lv, M. J., Zhu, X. G., and Chin, F. Y. L. (2013). IDBA-tran: A more robust de novo de Bruijn graph assembler for transcriptomes with uneven expression levels. Bioinformatics 29, 326–334. doi:10.1093/bioinformatics/btt219.

Petti, C. A., Polage, C. R., and Schreckenberger, P. (2005). The Role of 16S rRNA Gene Sequencing in Identification of.pdf. J. Clin. Microbiol. 43, 6123–6125. doi:10.1128/JCM.43.12.6123.

Pettorelli, N., Graham, N. A. J., Seddon, N., Maria da Cunha Bustamante, M., Lowton, M. J., Sutherland, W. J., et al. (2021). Time to integrate global climate change and biodiversity science-policy agendas. J. Appl. Ecol. 58, 2384–2393. doi:10.1111/1365-2664.13985.

Piper, A. M., Batovska, J., Cogan, N. O. I., Weiss, J., Cunningham, J. P., Rodoni, B. C., et al. (2019). Prospects and challenges of implementing DNA metabarcoding for high-throughput insect surveillance. Gigascience 8, 1–22. doi:10.1093/gigascience/giz092.

Quast, C., Pruesse, E., Yilmaz, P., Gerken, J., Schweer, T., Yarza, P., et al. (2013). The SILVA ribosomal RNA gene database project: Improved data processing and web-based tools. Nucleic Acids Res. 41, 590–596. doi:10.1093/nar/gks1219.

Quince, C., Walker, A. W., Simpson, J. T., Loman, N. J., and Segata, N. (2017). Shotgun metagenomics, from sampling to analysis. Nat. Biotechnol. 35, 833–844. doi:10.1038/nbt.3935.

Reback, J., McKinney, W., jbrockmendel, Bossche, J. Van den, Augspurger, T., Cloud, P., et al. (2020). pandas-dev/pandas: Pandas 1.0.3. doi:10.5281/ZENODO.3715232.

Reid, A. J., Carlson, A. K., Creed, I. F., Eliason, E. J., Gell, P. A., Johnson, P. T. J., et al. (2019). Emerging threats and persistent conservation challenges for freshwater biodiversity. Biol. Rev. 94, 849–873. doi:10.1111/brv.12480.

Resh, V. H., and Unzicker, J. D. (1975). Water Quality Monitoring and Aquatic Organisms : The Importance of Species Identification. Water Pollut. Control Fed. 47, 9–19.

Robertson, G., Schein, J., Chiu, R., Corbett, R., Field, M., Jackman, S. D., et al. (2010). De novo assembly and analysis of RNA-seq data. Nat. Methods 7, 909–912. doi:10.1038/nmeth.1517.

Sagova-Mareckova, M., Boenigk, J., Bouchez, A., Cermakova, K., Chonova, T., Cordier, T., et al. (2021). Expanding ecological assessment by integrating microorganisms into routine freshwater biomonitoring. Water Res. 191, 116767. doi:10.1016/j.watres.2020.116767.

scikit-bio development team, T. (2020). scikit-bio: A Bioinformatics Library for Data Scientists, Students, and Developers. Available at: http://scikit-bio.org.

Seabold, S., and Perktold, J. (2010). Statsmodels: Econometric and Statistical Modeling with Python. Proc. 9th Python Sci. Conf., 92–96. doi:10.25080/majora-92bf1922-011.

Seemann, T. BAsic Rapid Ribosomal RNA Predictor - barrnap. Available at: https://github.com/tseemann/barrnap.

Shakya, M., Lo, C. C., and Chain, P. S. G. (2019). Advances and challenges in metatranscriptomic analysis. Front. Genet. 10, 1–10. doi:10.3389/fgene.2019.00904.

Shi, Y., Tyson, G. W., Eppley, J. M., and Delong, E. F. (2011). Integrated metatranscriptomic and metagenomic analyses of stratified microbial assemblages in the open ocean. ISME J. 5, 999–1013. doi:10.1038/ismej.2010.189.

Sierra, M. A., Li, Q., Pushalkar, S., Paul, B., Sandoval, T. A., Kamer, A. R., et al. (2020). The Influences of Bioinformatics Tools and Reference Databases in Analyzing the Human Oral Microbial Community. Genes (Basel). 11. doi:10.3390/genes11080878.

Smith, M. B., Rocha, A. M., Smillie, C. S., Olesen, S. W., Paradis, C., Wu, L., et al. (2015). Natural Bacterial Communities Serve as Quantitative Geochemical Biosensors. MBio 6, e00326–15. doi:10.1128/mBio.00326-15.

Stat, M., Huggett, M. J., Bernasconi, R., Dibattista, J. D., Berry, T. E., Newman, S. J., et al. (2017). Ecosystem biomonitoring with eDNA: Metabarcoding across the tree of life in a tropical marine environment. Sci. Rep. 7, 1–11. doi:10.1038/s41598-017-12501-5.

Stein, E. D., White, B. P., Mazor, R. D., Jackson, J. K., Battle, J. M., Miller, P. E., et al. (2014). Does DNA barcoding improve performance of traditional stream bioassessment metrics? Freshw. Sci. 33, 302–311. doi:10.1086/674782.

Stoeck, T., Kochems, R., Forster, D., Lejzerowicz, F., and Pawlowski, J. (2018). Metabarcoding of benthic ciliate communities shows high potential for environmental monitoring in salmon aquaculture. Ecol. Indic. 85, 153–164. doi:10.1016/j.ecolind.2017.10.041.

Sweeney, B. W., Battle, J. M., Jackson, J. K., and Dapkey, T. (2011). Can DNA barcodes of stream macroinvertebrates improve descriptions of community structure and water quality? J. North Am. Benthol. Soc. 30, 195–216. doi:10.1899/10-016.1.

Taberlet, P., Coissac, E., Pompanon, F., Brochmann, C., and Willerslev, E. (2012). Towards next-generation biodiversity assessment using DNA metabarcoding. Mol. Ecol. 21, 2045–2050. Available at: http://onlinelibrary.wiley.com/doi/10.1111/j.1365-294X.2012.05470.x/full%5Cnpapers2://publication/uuid/30F6E470-48F9-4C24-A3C1-6964EE26B34F.

Torti, A., Lever, M. A., and Jørgensen, B. B. (2015). Origin, dynamics, and implications of extracellular DNA pools in marine sediments. Mar. Genomics 24, 185–196. doi:10.1016/j.margen.2015.08.007.

Turner, T. R., Ramakrishnan, K., Walshaw, J., Heavens, D., Alston, M., Swarbreck, D., et al. (2013). Comparative metatranscriptomics reveals kingdom level changes in the rhizosphere microbiome of plants. ISME J. 7, 2248–2258. doi:10.1038/ismej.2013.119.

Urich, T., Lanzén, A., Qi, J., Huson, D. H., Schleper, C., and Schuster, S. C. (2008). Simultaneous assessment of soil microbial community structure and function through analysis of the meta-transcriptome. PLoS One 3. doi:10.1371/journal.pone.0002527.

Urich, T., Lanzén, A., Stokke, R., Pedersen, R. B., Bayer, C., Thorseth, I. H., et al. (2014). Microbial community structure and functioning in marine sediments associated with diffuse hydrothermal venting assessed by integrated meta-omics. Environ. Microbiol. 16, 2699– 2710. doi:10.1111/1462-2920.12283.

van der Walt, A. J., van Goethem, M. W., Ramond, J. B., Makhalanyane, T. P., Reva, O., and Cowan, D. A. (2017). Assembling metagenomes, one community at a time. BMC Genomics 18, 1–13. doi:10.1186/s12864-017-3918-9.

Van Rossum, G., and Drake, F. L. (2009). Python 3 Reference Manual. Scotts Valley, CA: CreateSpace.

Virtanen, P., Gommers, R., Oliphant, T. E., Haberland, M., Reddy, T., Cournapeau, D., et al. (2020). SciPy 1.0: fundamental algorithms for scientific computing in Python. Nat. Methods 17, 261–272. doi:10.1038/s41592-019-0686-2.

Vollmers, J., Wiegand, S., and Kaster, A. K. (2017). Comparing and evaluating metagenome assembly tools from a microbiologist’s perspective - Not only size matters! PLoS One 12, 1– 31. doi:10.1371/journal.pone.0169662.

Wang, Y., Hu, H., and Li, X. (2017). rRNAFilter: A Fast Approach for Ribosomal RNA Read Removal Without a Reference Database. J. Comput. Biol. 24, 368–375. doi:10.1089/cmb.2016.0113.

Westermann, A. J., Gorski, S. A., and Vogel, J. (2012). Dual RNA-seq of pathogen and host. Nat. Rev. Microbiol. 10, 618–630. doi:10.1038/nrmicro2852.

Will, K. W., and Rubinoff, D. (2004). Myth of the molecule: DNA barcodes for species cannot replace morphology for identification and classification. Cladistics 20, 47–55. doi:10.1111/j.1096-0031.2003.00008.x.

Wood, D. E., Lu, J., and Langmead, B. (2019). Improved metagenomic analysis with Kraken 2. Genome Biol. 20, 1–13. doi:10.1186/s13059-019-1891-0.

Wooley, J. C., Godzik, A., and Friedberg, I. (2010). A primer on metagenomics. PLoS Comput. Biol. 6. doi:10.1371/journal.pcbi.1000667.

WWF (2020). Living Planet Report 2020 - Bending the curve of biodiversity loss., eds. R. E. A. Almond, M. Grooten, and T. Petersen Gland, Switzerland.

Yan, Y. W., Jiang, Q. Y., Wang, J. G., Zhu, T., Zou, B., Qiu, Q. F., et al. (2018). Microbial communities and diversities in mudflat sediments analyzed using a modified metatranscriptomic method. Front. Microbiol. 9, 1–15. doi:10.3389/fmicb.2018.00093.

Yates, M. C., Derry, A. M., and Cristescu, M. E. (2021). Environmental RNA: A Revolution in Ecological Resolution? Trends Ecol. Evol. 36, 601–609. doi:10.1016/j.tree.2021.03.001.

Yilmaz, P., Kottmann, R., Field, D., Knight, R., Cole, J. R., Amaral-Zettler, L., et al. (2011). Minimum information about a marker gene sequence (MIMARKS) and minimum information about any (x) sequence (MIxS) specifications. Nat. Biotechnol. 29, 415–420. doi:10.1038/nbt.1823.

Yu, K., and Zhang, T. (2012). Metagenomic and metatranscriptomic analysis of microbial community structure and gene expression of activated sludge. PLoS One 7. doi:10.1371/journal.pone.0038183.

