## Supplementary material for "Metagenomics vs. total RNA sequencing: most accurate data-processing tools, microbial identification accuracy, and implications for freshwater assessments"

### SUPPLEMENTAL MATERIAL

#### Supplemental material 1: processing steps and quality control of DNA and RNA library preparation and shotgun sequencing, obtained from the sequencing center Génome Québec

Total RNA was quantified using a NanoDrop Spectrophotometer ND-1000 (NanoDrop Technologies, Inc.) (Tab. S1), and its integrity was assessed on a 2100 Bioanalyzer (Agilent Technologies) (Tab S3). Since the samples contained both prokaryotic and eukaryotic RNA, RINs were not applicable, and low RINs were ignored. Libraries were generated from 1 µL of each sample as follows: cDNA synthesis was achieved with the NEBNext RNA First Strand Synthesis E7771 and NEBNext Ultra Directional RNA Second Strand Synthesis Modules (New England Biolabs; Whitby; ON Canada). The remaining steps of library preparation were done using the NEBNext Ultra II DNA Library Prep Kit for Illumina (New England Biolabs; Whitby; ON Canada). Adapters and PCR primers were purchased from New England Biolabs (Whitby; ON Canada). Libraries were quantified using the Kapa Illumina GA with Revised Primers-SYBR Fast Universal kit (Roche Sequencing Solutions Inc; Pleasanton; CA U.S.A). The average fragment size was determined using a LabChip GXII instrument (PerkinElmer). Note that the mRNA enrichment step was skipped to create total RNA libraries.

gDNA was quantified using the Quant-iT™ PicoGreen® dsDNA Assay Kit (Thermo Fisher Scientific; Burlington; ON Canada) (Tab. S2). Libraries were generated using the NEBNext Ultra II DNA Library Prep Kit for Illumina (New England Biolabs; Whitby; ON Canada) as per the manufacturer's recommendations. Adapters and PCR primers were purchased from IDT (Coralville; IA U.S.A.). Size selection of libraries for the desired insert size was performed using SparQ beads (VWR; Mississauga; ON Canada). Libraries were quantified using the Kapa Illumina GA with Revised Primers-SYBR Fast Universal kit (Roche Sequencing Solutions Inc; Pleasanton; CA U.S.A). The average fragment size was determined using a LabChip GXII instrument (PerkinElmer). 5 µL of both the DNA and RNA libraries were respectively combined and used for quality control.

During library preparation, normalization was performed by processing equal volumes of samples instead of equal concentrations of samples. We chose this alternative normalization method because it allowed for an equal relative sequencing depth per sample as opposed to an equal total sequencing depth. That way, the relative numbers of reads per sample mirrored the relative amount of DNA/RNA in each sample, avoiding an over- or underrepresentation of samples with higher or lower DNA/RNA amounts. The DNA libraries yielded fragments of approximately 488 bp length, whereas the RNA libraries yielded fragments of approximately 295 bp length (both including adaptors and indices). For sequencing, the DNA shotgun library pools and the total RNA library pools were mixed according to their concentration so that each represented 50% of the data. The libraries were first normalized at 2 nM and then pooled and denatured in 0.04 N NaOH. The pool was diluted to 12 pM using HT1 buffer, loaded onto a MiSeq, and sequenced for 2x150 cycles according to the manufacturer's instructions. The MiSeq Control Software (MCS) version was 2.5.0.5 and the Real-Time Analysis (RTA) version was 1.18.54. The program bcl2fastq v1.8.4 was then used to demultiplex samples and to generate fastq reads.

#### Supplemental material 2: setup of BLAST and kraken2 databases for SILVA

We downloaded the available SILVA132_NR99 SSU and LSU databases and merged them into one SILVA database, removing duplicates that falsely occurred in both databases according to the SILVA-arb support. Due to our initially planned comparison between SILVA and Genbank, we standardized the taxonomy nomenclature of the SILVA database by translating the taxonomy of all SILVA reference sequences into the Genbank taxonomy before building SILVA reference databases. Note that some SILVA reference sequences are annotated with species names that do not match the rest of their taxonomic annotation. To translate the SILVA taxonomy into Genbank taxonomy, we first made sure that the last taxonomic rank of each SILVA reference sequence was a species name that fit the remaining taxonomy. Therefore, we checked if the genus of the species name matched the genus name for each SILVA reference sequence, and if not, the species name was converted to *NA*. Then, we checked each taxonomic rank of each SILVA reference sequence for matches among first scientific and then non-scientific names in the Genbank taxonomy file *names.dmp (*available through the NCBI archive, part of *taxdmp.zip*, https://ftp.ncbi.nlm.nih.gov/pub/taxonomy/), beginning at the last rank, and if a match was found, the respective Genbank taxonomic ID was used for that SILVA reference sequence. If no match was found for that rank, the process was repeated for the next higher rank until a match was found. If no match was found overall, the reference sequence was assigned with *NA*. Note that we removed all Genbank names containing “environmental”, “uncultured”, “unidentified”, and “metagenome” from the *names.dmp* file prior to translation to exclude them from the matching process.

kraken2 databases were set up following the developer’s instructions (https://github.com/DerrickWood/kraken2/blob/master/docs/MANUAL.markdown). To set up a kraken2 database for the merged SSU and LSU SILVA database, we translated the taxonomy of each reference sequence into Genbank taxonomy IDs as described above, added the IDs to each reference sequence, and created files in the same format as the taxonomy files of the Genbank database, *names.dmp* and *nodes.dmp* (part of *taxdmp.zip*, https://ftp.ncbi.nlm.nih.gov/pub/taxonomy/). We automated that process with a script that is available on GitHub (https://github.com/hempelc/metagenomics-vs-totalRNASeq), which incorporates the SILVA taxonomy files *taxmap_slv_ssu_ref_nr_138.1.txt.gz*, *taxmap_slv_lsu_ref_nr_138.1.txt.gz*, and *tax_slv_ssu_138.1.txt.gz* available through the SILVA archive (https://www.arb-silva.de/download/archive/).

BLAST databases were set up using the makeblastdb command with the parameter `--taxid_map` to incorporate taxonomic IDs, including translated IDs for SILVA reference sequences, into the respective BLAST databases.

#### Supplemental tables

*Table S1: Nanodrop Quantification (RNA). 1-3=replicates; ExtCon=extraction control; FilCon=filtration control.*

| Sample | Concentration [ng/µl] | Total RNA [ng] | 260/230 | 260/280 |
|---|---|---|---|---|
| RNA 1 | 1.87 | 33.66 | 0.36 | 8.19 |
| RNA 2 | 3.32 | 59.76 | 0.39 | 2.02 |
| RNA 3 | 2.96 | 53.28 | 0.33 | 3.79 |
| RNA FilCon | 0.73 | 13.14 | 0.09 | -1.43 |
| RNA ExtCon | 0.43 | 7.74 | 0.08 | 4.22 |

*Table S2: Fluorescence Assay Quantification (DNA). 1-3=replicates; ExtCon=extraction control; FilCon=filtration control.*

| Sample | Concentration [ng/µl] | Total DNA [ng] |
|---|---|---|
| DNA 1 | 2.7856 | 136.494 |
| DNA 2 | 3.0507 | 149.484 |
| DNA 3 | 2.3648 | 115.875 |
| DNA FilCon | 0 | 0 |
| DNA ExtCon | 0 | 0 |

*Table S3: Bioanalysis (RNA). 1-3=replicates; ExtCon=extraction control; FilCon=filtration control.*

| Sample | 28S/18S | RIN | Concentration [ng/µl] | Total RNA [ng] |
|---|---|---|---|---|
| RNA 1 | 1.279624 | N/A | 1.87 | 33.66 |
| RNA 2 | 1.1748 | N/A | 3.32 | 59.76 |
| RNA 3 | 1.123146 | N/A | 2.96 | 53.28 |
| RNA FilCon | 0 | 1.2 | 0.73 | 13.14 |
| RNA ExtCon | 0 | 1.7 | 0.43 | 7.74 |

#### Supplemental Figures

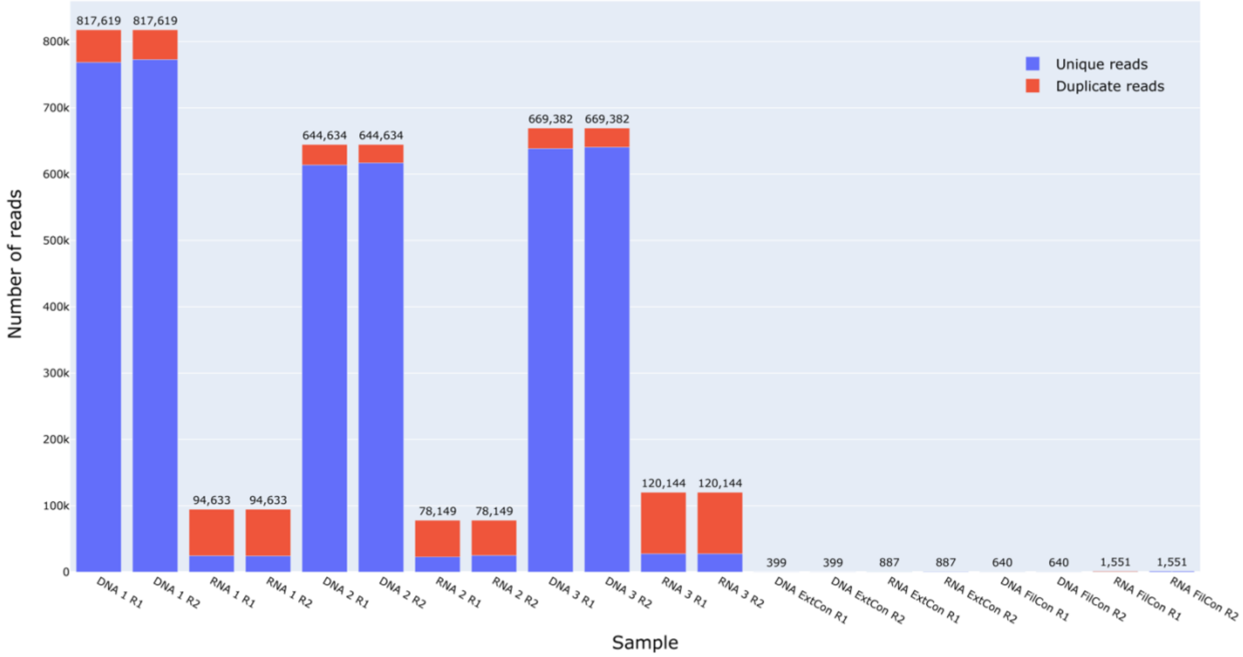


*Figure S1: Number of reads per sample.*

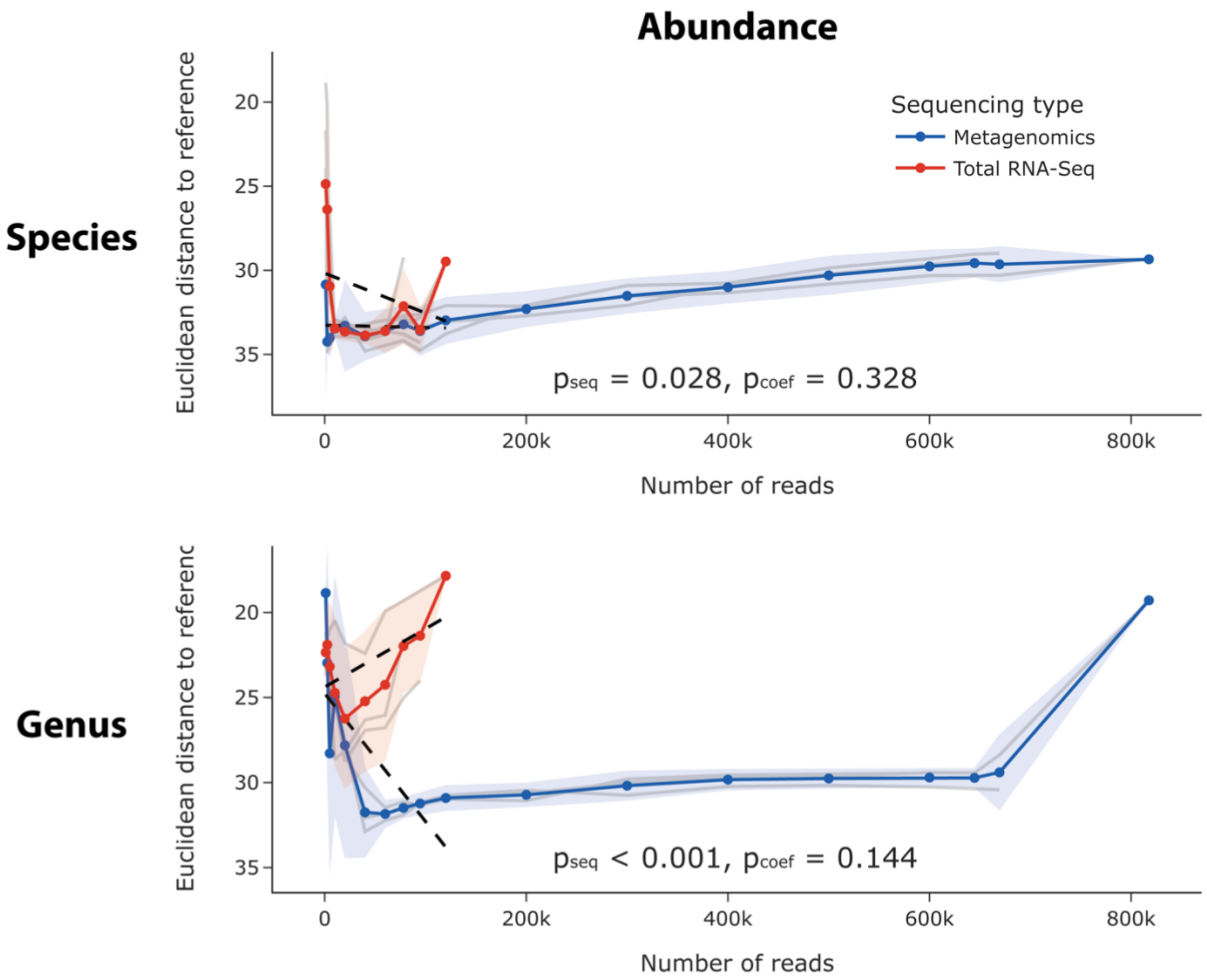


*Figure S2: Relationship between sequencing depth and accuracy for abundance-based evaluation levels, including subsamples up to 20,000 reads. Evaluation levels consisted of combinations of data type (abundance/P-A), and taxonomic rank (genus/species). Blue and red lines indicate the mean Euclidean distance of all metagenomics and total RNA-Seq replicates, which have each been subsampled ten times, and the area around the lines indicates the standard deviation (SD). Lower Euclidean distances are a proxy for higher accuracy. The y-axis is inverted, and its scale varies among graphs. The SD equals zero at the highest number of reads since all available reads were used and, therefore, no subsamples could be generated. Individual replicates are shown as grey lines. Regression curves are shown as dashed black lines for the portion of the data that was comparable between metagenomics and total RNA-Seq. $p_{seq}$-values are based on partial F-tests between linear models including or excluding the sequencing method (metagenomics/total RNA-Seq) as a binary independent variable. $p_{coef}$-values are based on paired t-tests between the coefficients of regression curves of individual metagenomics and total RNA-Seq replicates based on the comparable portion of the data.*

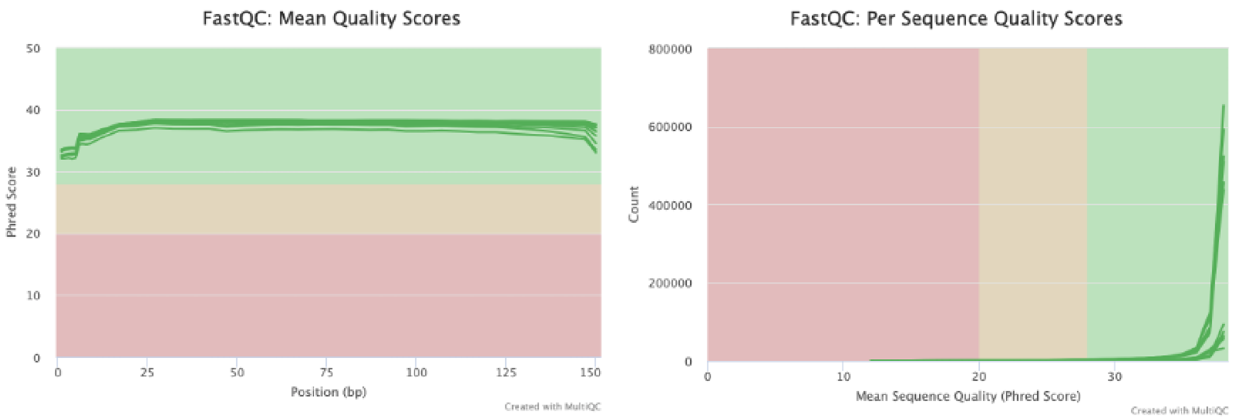


*Figure S3: Mean and per sequence quality scores.*
